# Assessing above and belowground recovery from ammonium sulfate addition and wildfire in a lowland heath: mycorrhizal fungi as potential indicators

**DOI:** 10.1101/2023.07.01.547153

**Authors:** Jill Kowal, Raquel Pino-Bodas, Elena Arrigoni, Guillaume Delhaye, Laura M. Suz, Jeffrey G. Duckett, Martin I. Bidartondo, Silvia Pressel

**Author notes:** **Corresponding Author:** Dr. Jill Kowal. **Author Contributions** JK, RP, conceived the ideas and designed methodology; all authors but GD collected field and laboratory data; all authors analysed the data; JK led the writing of the manuscript. All authors contributed critically to the drafts and gave final approval for publication.

## Abstract

Atmospheric pollution containing soil-nitrifying ammonium sulphate is affecting semi-natural ecosystems worldwide. Long-term additions of ammonium sulphate on nitrogen (N)-limited habitats such as heathlands increase climate stress affecting recovery from wildfires. Yet although heathland vegetation largely depends on ericoid mycorrhizal fungi (ErM) to access soil N, we lack a detailed understanding of how prolonged exposure to ammonium sulphate may alter ErM community composition and host plants’ reliance on fungal partners following wildfire and substantial reductions in ammonium sulphate pollution. Effects on ecosystem processes, particularly carbon stores, also remain uncertain. Ammonium sulphate additions occurred bi-weekly for five years after a 2006 wildfire burnt a UK heathland. Ten years after the treatments ceased (2021), we measured vegetation structure, lichen and lichen photobiont composition, soil characteristics, ErM colonisation, ErM diversity in roots and soil, and assessed their potential as recovery indicators. We found heather height and density, and moss groundcover, were greater in N-enriched plots. Lichen community indices showed significant treatment effects without photobiont differences. Soil pH and Mg, and the proportion of putative ErM fungi in soil were significantly lower in treated plots while soil cation exchange capacity was significantly higher. Increases in soil pH were positively correlated to soil ErM abundance. Soil carbon stock measures were variable and negatively related with soil ErM. Our results indicate that atmospheric pollution following fire can have significant lingering effects and mycorrhizal fungi diversity are a novel and effective ecological tool to assess ecosystem recovery on heathlands.

**IMPLICATIONS FOR PRACTICE:**

- Recovery of heathlands from wildfire and atmospheric pollution may require decadal scales. Conventional restoration assessment tools lack critical understanding of belowground soil and mycorrhizal fungi interactions and nutritional feedback loops with aboveground hosts.
- Given that ericoid mycorrhizal fungi (ErM) communities are key nutrient regulators of resilient heathlands, we recommend baseline measurements of ErM diversity and abundance in both soil and dominant plant roots be added to the recovery assessment toolkit prior to commencing restoration or management plans.
- What defines ErM recovery, and disentangling effects of ammonium sulphate pollution and fire, remain open questions which only long-term field experiments across pollution gradients will address.

## Introduction

Plants depend largely on their mycorrhizal fungal symbionts to access soil nitrogen (N) and phosphorus (P) particularly in nutrient-poor heathlands (Read 1996; Smith & Read 2008). Questions relating to the impact of atmospheric pollution on mycorrhizal fungal communities and their function in N- and P-limited systems such as heathlands are becoming more pressing as the ecosystem services that heathlands provide, especially regarding carbon (C) sequestration, come into focus. Particularly critical are the contributions of the dominant *Calluna vulgaris* roots and associated ericoid mycorrhizal fungi (ErM) to belowground C (Clemmensen et al. 2013) and how these may be affected by N deposition (Van Geel et. al. 2020), given the key role of ErM in plant N acquisition (Leake et al. 1990; van der Heijden et al. 2015). Nitrogen-limited heathlands across Europe are particularly vulnerable to atmospheric N deposition (Southon et al. 2013; Van Geel et al. 2020) which threatens their ecosystem function through loss of vegetation biodiversity aboveground (Tang et al. 2018; van Paassen et al. 2020) and soil changes affecting microbial communities belowground (Treseder 2008; Zhou et al. 2017; Kowal et al. 2022). Long-term N addition has been shown to alter root-associated fungal community composition, increase ErM root colonisation (Kiheri et al. 2020) or alleviate plant (heather) N deficits, thereby reducing reliance on ErM (Vesala et al. 2021). *Calluna vulgaris* age also plays a role in ecosystem function (Field et al. 2017).

European dry heathlands are reservoirs of biodiversity and provide essential ecosystem services such as C sequestration (Friggens et al. 2020; Gregg et al. 2021), climate regulation and flood mitigation (Alonso et al. 2012). Currently, heathlands are threatened by anthropogenic pressures including encroachment by trees and agriculture plus soil acidification and eutrophication from transboundary pollutants, especially ammonium sulphate, a common N fertilizer and key component of particulate matter (PM) (EMEP 2019; Chen et al. 2018). Dry heathlands are dominated by dwarf ericaceous shrubs which associate with ErM and support diverse lichen and bryophyte communities. Whether heathlands and their belowground ErM can recover following substantial reductions in ammonium sulphate is difficult to assess on a field-scale.

Appreciation of the diversity of putative ErM colonising Ericaceae roots has expanded recently (Bougoure et al., 2007; Smith & Read 2008, Fehrer et al. 2019) to include many ascomycetes in the order Helotiales (Walker et al., 2011) and some Basidiomycota (Selosse et al. 2007; Vohnick et al. 2012). It is common for multiple ErM species to co-colonise a single host plant (Gorzelak et al. 2012) and their functionality, when combined, might become enhanced leading to increased resilience to environmental change (Perotto et al. 2002). Van Geel et al. (2020) identified 239 putative ErM operational taxonomic units (OTUs) associating with heather roots alone across 21 dry heathlands. Another recent study of ErM diversity in Norwegian heathlands found several new ErM OTUs within both Ascomycota and Basidiomycota associating with heather roots (Blaalid & Davey 2022). Further ErM diversity is being uncovered using different DNA sequence clustering methods, such as amplicon sequence variants (ASVs), which can now be linked to species hypotheses (SH) (Kõljalg et al. 2020; Põlme et al. 2020; Abarenkov et al. 2021).

Lichens are sensitive to air quality and have long been used as bioindicators (Nash 1996; Nimis et al. 2002; Giordani 2007). However, the effects of N deposition on terricolous lichen communities, typical of heathlands, have been largely ignored with only a few studies to date showing that N deposition negatively affected some heathland lichen species (Pilkington et al. 2007; Stevens et al. 2012; Edmondson et al. 2013). This represents a considerable knowledge gap since terricolous lichens provide important ecosystem services including soil stabilization, soil moisture retention and facilitation of seed germination (Asplund & Wardle 2017). Moreover, few studies have investigated the recovery of lichen communities once disturbance ceased (Edmondson et al. 2013; Benvenutto-Vargas & Ochoa-Hueso 2020). While variation in the composition of lichen communities under atmospheric pollution and different climatic gradients is well documented (Gadsdon et al. 2010), little is known about potential changes in the interactions between mycobiont and photobiont; this is surprising since adaptation of lichens to different environmental conditions is highly dependent on their photobiont (Peksa & Škaloud 2011). The ability of mycobionts to establish associations with several lineages of photobionts is considered an adaptive strategy to tolerate a wider range of environmental conditions (Fernandez-Mendoza et al. 2011; Jüriado et al. 2019). Recent studies show that photobionts switch along environmental gradients (Dal Grande et al. 2018); however, the effect of N deposition on mycobiont-photobiont interactions has not been investigated.

In this study we examined a subset of the experimental plots used by Southon et al. (2012) to determine whether there are lingering effects on ecosystem function more than ten years since the last ammonium sulphate addition and wildfire (for a description of the Southon et al.’s experiments, see Background Data S1, Table S1). Specifically, we asked whether differences between the N-treated and control plots were still detectable (1) aboveground in the structure or diversity of plant and lichen communities and associated photobionts, and (2) belowground in soil C stocks and mycorrhizal and soil fungal community composition. We propose that the slow growth of the dominant heather shrubs renders this habitat resistant to full recovery of vegetation structure and lichen community composition from ammonium sulphate addition. However, we expect recovery in the soil chemistry, soil fungi and mycorrhizal community composition given more than 10 years of precipitation and background pollution to diffuse treatment effects. Finally, we present baseline molecular data on the ErM fungal community to assess its potential as a heathland recovery metric.

## Methods

### Experimental design and site

The study was carried out in June 2021 at Thursley NNR in southern England (51°09’25” N 0°42’04” W). The site comprises H2 dry heathland (National Vegetation Classification (NVC) scheme, Rodwell 2006) and is designated a Special Area of Conservation (European Union’s Habitats Directive, 92/43/EEC). The soil is a podzol derived from Lower Greensand with a thin (<1 cm) humus layer overlying a sandy mineral layer (Power et al., 1998). In 2021, the mean annual precipitation was 781 mm and mean annual temperature 11.9° C (minimum 6° C in January and maximum 20° C in August) (Fig. S1). Background N and sulphur (S) deposition levels were 14 kg ha^-1^ year^-1^ and 12.7 kg ha^-1^ year^-1^ in 2020 (www.worldweatheronline.com, Levy et al., 2020), higher than the N levels measured at the site (8 kg ha^-1^ year^-1^) from 1998 and 2000 (Power & Barker 2003); S alone was not measured.

We studied three of the four blocks assessed in Southon et al. (2012) (Fig. S2a, b, c); each block contains four 4 m^2^ plots, each with four 2 m^2^ subplots. These three blocks were selected as they were the only blocks with the same number of treatments; two control plots and two treated plots which received bi-weekly additions of ammonium sulphate at a rate of 30 kg^-1^ ha (dissolved and distributed using a fine nozzle spray) during 1998 through 2010. This rate approximated levels reported in Europe, e.g., the Netherlands, during this period (Bobbink et al. 2013).

### Vegetation and lichen survey

We carried out a vegetation and lichen survey of the three blocks. For each subplot we measured vegetation density, abundance and height, and recorded plant and lichen species composition. Following the lichen survey, three specimens of each lichen species were collected to confirm the identifications and carry out DNA identification of photobionts in the laboratory (Methods S1). Nomenclature for bryophytes follows Blockeel et al. (2021) and for lichens Burgaz et al. (2020) and Pino-Bodas et al. (2021).

### Soil sampling

To assess soil characteristics and chemistry, we collected two soil samples per plot to a depth of 15 cm. Each sample comprised three subsamples which combined gave at least 500 g dry weight (DW). Soil samples were stored at 4°C and processed by NRM Laboratories (Bracknell, England, Methods S2). We measured the following: water content, bulk density, dry matter, pH, total C and N concentrations (% g/100g, on a dry matter basis); N, P, potassium (K), magnesium (Mg), nitrate (NO_3_), ammonium (NH ^+^), sodium (Na), calcium (Ca), all (mg/kg); cation exchange capacity (CEC) and C stock (t/ha). Approximately 50 g of fresh soil was removed from each soil sample (taking care to remove at least 15 g from each subsample) and placed at -80°C for environmental DNA analysis (see below).

### Plant leaves, roots and stems analyses

*Calluna vulgaris* leaf tissue (50 g DW) was collected from at least three randomly selected plants from each plot (n=12) to measure foliar N, P and C. Immediately after collection, samples were dried (overnight at 60°C) and processed by NRM Laboratories (Bracknell, England; see Methods S2).

For ErM fungal root colonisation and community assessments, three young plants of *C. vulgaris* were collected. For each of their roots, one half was stored in ethanol (70% v/v) at 4°C for ink staining to assess ErM root colonisation, the other half was stored in cetyltrimethylammonium bromide (CTAB) extraction buffer at -20°C for molecular identification. Ericoid mycorrhizal colonisation was scored as present only when typical ErM hyphal coils were visible within the cell walls of at least three root cells per root section (Kowal et al. 2020a, b).

To assess lateral stem seasonal growth, we selected five mature plants of *C. vulgaris* per plot and removed the largest stem at the plant’s base. The stem cuttings were kept at 4°C for further processing and analysis. Stem cuttings were hand sectioned (approximately 0.25 mm thick) and annual growth ring widths were measured (x 3 per section) on both the widest and narrowest radial hemispheres (x40) under a Zeiss Axioscop 2 microscope equipped with an AxioCam MRc digital camera and auto-calibrated using Axiovision Microscope Software (Fig. S3). The results were tabulated and aligned to compare years of treatment and the years post-treatment.

### Soil fungal community analysis

Total soil DNA was extracted in duplicate from 0.25 g of soil from each plot (12 plots, 24 extractions, DNeasy PowerSoil Pro Kit, Qiagen, Hilden, Germany). DNA extraction quality (260/280nm and 260/230nm ratios) was assessed using a NanoDrop2000 spectrophotometer (Thermo Fisher Scientific) and DNA concentration was quantified using a Quantus^™^ Fluorometer and QuantiFluor® dsDNA System (Promega, Madison, Wisconsin, United States). The DNA extractions were processed by Macrogen (Seoul, South Korea) for library preparation and Illumina sequencing with the primer set ITS86F and ITS4 (White et al. 1990; Turenne et al. 1999) targeting the ITS2 region.

### Root fungal community analysis

To identify the ErM fungal communities colonising *C. vulgaris* roots, we pooled ten roots from each of the heather plants sampled per plot (30 roots) and carried out three DNA extractions (DNeasy PowerSoil Pro Kit, Qiagen, Hilden, Germany) for each plot (n = 12 extractions per block, 36 extractions overall) followed by amplification, cloning and sequencing of the complete ITS region (Methods S3). An ABI3730 genetic analyser (Applied Biosystems) was used for Sanger DNA sequencing, and forward and reverse sequences were assembled using Geneious v. 8.1.9. Partial SSU and LSU regions were trimmed using ITSx (Bengtsson-Palme et al. 2013) and the resulting full-length ITS sequences were clustered into Amplicon Sequence Variants (ASVs) in VSEARCH (Rognes et al., 2016). Amplicon Sequence Variants forming new Species Hypothesis (SHs) were excluded from further analyses as their taxonomic classification remains poorly resolved.

For both roots and soil, taxonomic classification was performed using the SH matching v2.0.0 tool on the PlutoF workbench (Abarenkov et al. 2010). Those assigned to the same SH were grouped and categorised by putative ecological function (mycorrhiza, ericoid associate or other) (Vohnik et al. 2012, 2016; Grelet et al. 2017; Van Geel et al. 2020). For statistical analysis, we only analysed taxa (OTUs) represented by > 100 reads. The overall proportion of ErM and non-ErM fungi was calculated as the total ErM and non-ErM sequences over the total number of sequences that passed the quality control in the SH matching analysis.

To compare our baseline analysis with a recent European heathland survey of root-derived ErM diversity, we grouped the OTUs reported in Van Geel et al. (2020) into SHs by filtering only OTUs derived from *C. vulgaris* roots in heathlands and assigning a putative ecological function to the SH taxonomy (Methods S3; see Methods S4 for the analysis of lichen photobiont DNA).

### Statistical analyses

Statistical analyses were conducted in R (R core Team 2022, v. 4.1.2) unless otherwise stated. Soil characteristics, leaf chemistry, aboveground biomass measures, lichen richness, abundance and diversity (Shannon and inverse Simpson indexes) were compared between treatments using linear mixed effects models, using block and replicates as random variables (function brm in the *brms* package). Shannon and inverse Simpson indexes were calculated using BiodiversityR (Kindt & Kindt 2019). Wilcoxon rank sum tests were applied with continuity correction. The effect of the treatment on the composition of total fungal and ErM fungal root and soil communities was tested using a nested PERMANOVA (nperm = 999) with block as a random variable on a Bray-Curtis dissimilarity matrix, using the *adonis2* function in the *vegan* package (Oksanen et al. 2013). The predicted values for both soil ErM richness and ErM colonisation rates were tested using soil covariates in simple linear regression models.

Effect of treatment on ErM colonisation presence or absence in heather roots was also tested using chi-square tests with Yates correction using Graphpad Prism (V 9.5 525). Correlation matrices were used to test ErM root colonisation and root and soil richness with soil chemical characteristics (Prism V 9.5 525).

To test significant differences between lichen community composition in control and N-addition subplots PERMANOVA (nperm = 999) was applied in *vegan*. The Bray-Curtis dissimilarity between communities was represented graphically by non-metric multidimensional scaling (NMDS) ordination. Indicspecies (De Cáceres & Jansen 2015) was used to identify lichen indicator species for each treatment, performed with *multpatt* (999 permutations).

## Results

### Vegetation and lichen structure, composition, abundance and diversity

The results of the vegetation and lichen surveys are summarized in Tables S2a, b, c. *Calluna vulgaris* dominated the shrub community and was present in every subplot. *Ulex europaeus and Erica tetralix* were present occasionally and with significantly less coverage with 112 and 24 grid points, respectively, compared with 2,429 *C. vulgaris* points. The ground layer consisted of either leaf litter, bryophytes (the mosses *Hypnum jutlandicum*, *Campylopous introflexus* and *Dicranum scoparium* and the leafy liverwort *Cephaloziella divaricata*) or a combination of the two along with 21 species of lichens, all belonging to the genus *Cladonia*. The most frequent *Cladonia* species were *C. furcata*, *C. crispata* subsp. *cetrariiformis*, *C. portentosa* and *C. verticillata*.

The mean height and tallest points of the heather canopy were both significantly higher in the N-addition plots than the control plots (ANOVA, *P* value <0.05) but no treatment differences were detected in the vegetation density at either ground level or 20cm (Fig. 1).

**Fig. 1.**
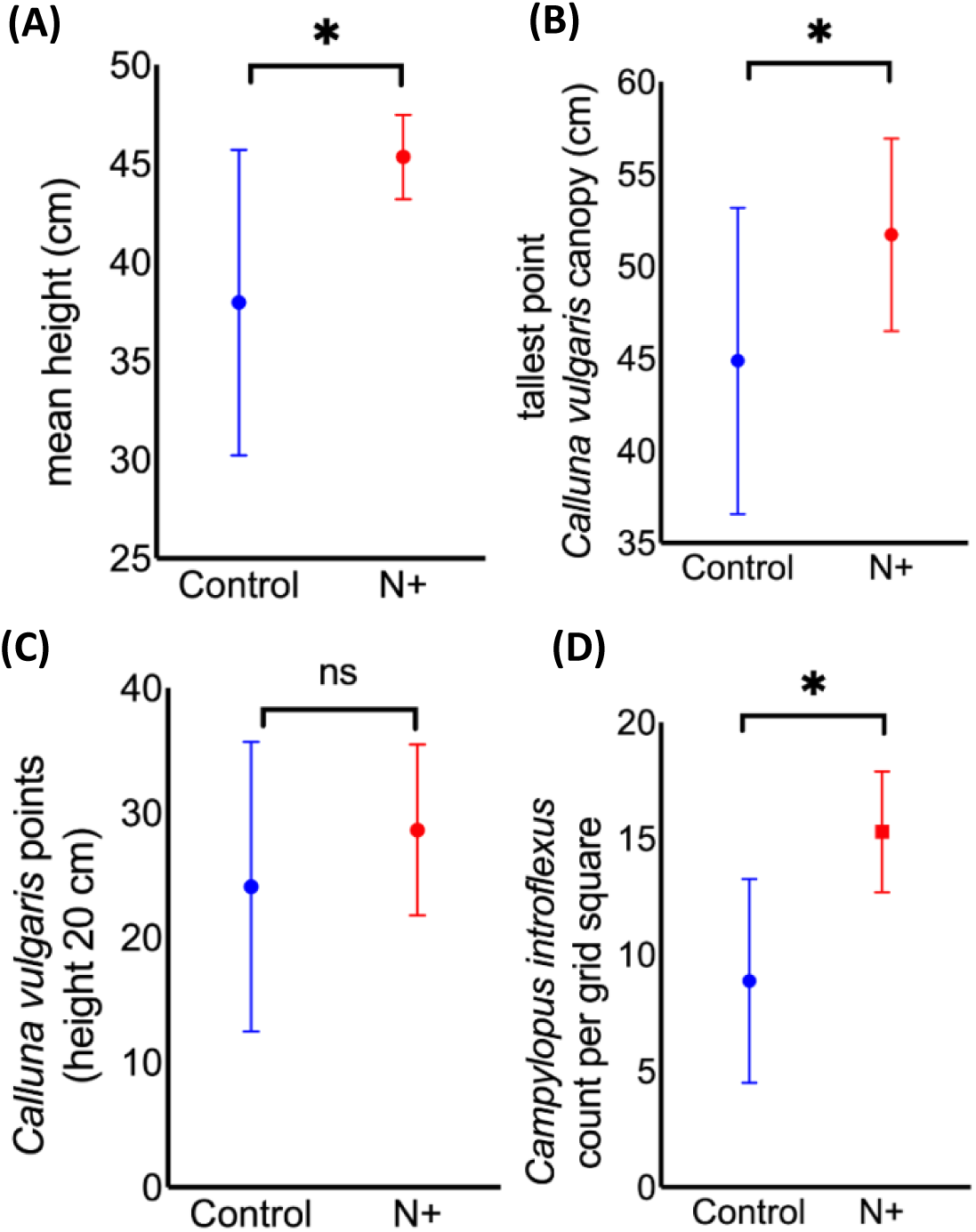
Comparison of aboveground biomass measures, between treatments (±SD). (A) mean height of vegetation; (B) tallest point within a subplot; (C) number of *Calluna vulgaris* points per quadrat (50 cm x 50 cm) counted at 20cm; and (D) *Campylopus introflexus* ground cover. Means ± SD.

*Campylopus introflexus* ground cover was significantly higher in the N-addition plots compared with the control plots (Table S3; Fig. 2D) but no treatment effects were seen in the other bryophyte species. Similarly, we found no differences in heather lateral stem growth between the two treatment plots when measuring growth rings in the immediate years following the fire (2006), during the years of further N-addition (2006-2011), or after N additions ceased (2012 to 2021 harvest) (Table S4). However, the fixed effect of year shows significant differences in growth patterns (*P* value: < 0.0001) with heather growth more robust in the first three years (average 0.36 mm/year) and slower in the last three years (0.16 mm/year). Notably, we observed a growth peak in the N-addition plots around 2014, two years after treatments ceased (Fig. S4).

**Fig. 2.**
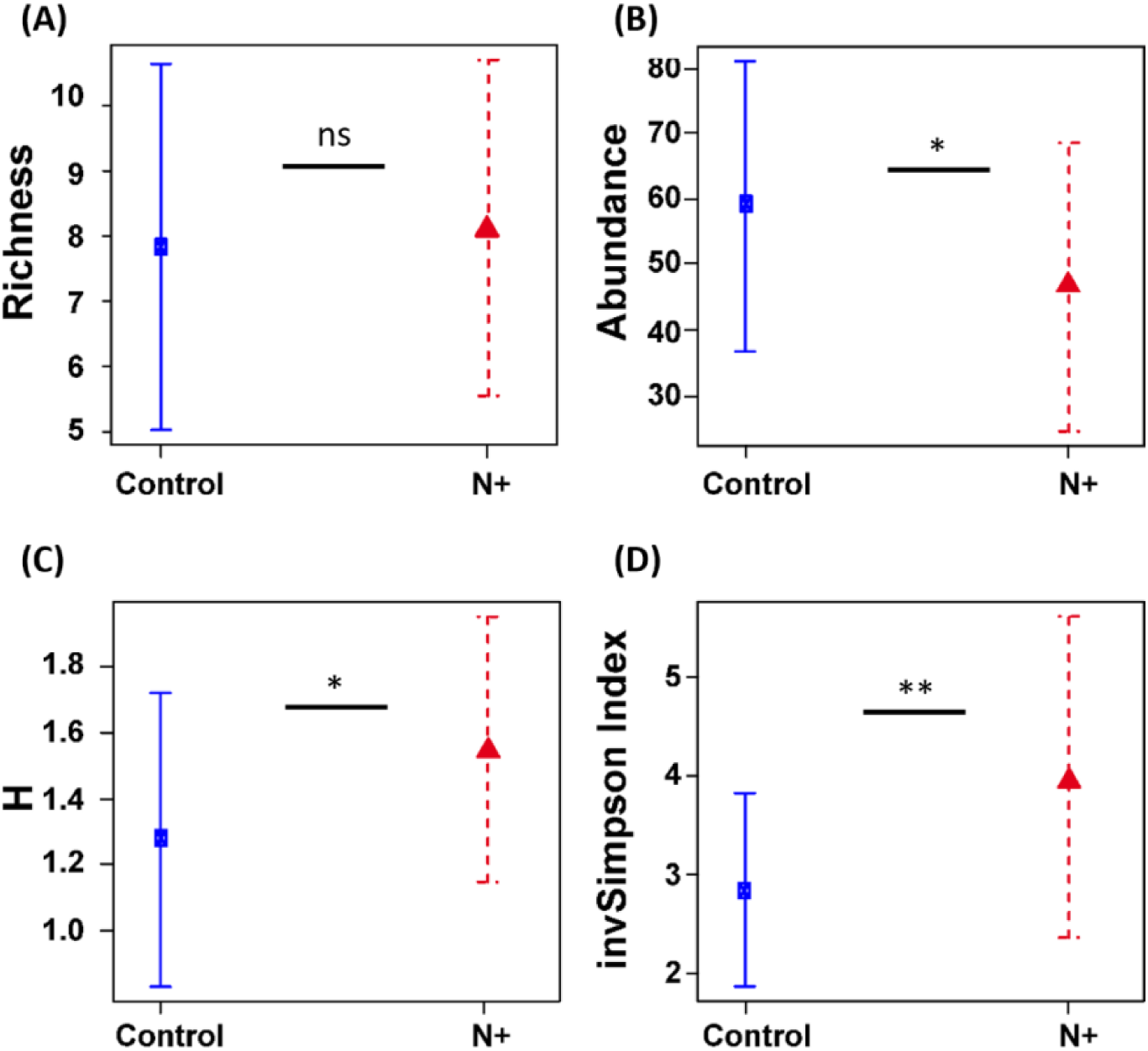
Treatment effects on lichen richness (A); abundance (B); diversity using Shannon Index, H (C); and, diversity, using inverse Simpson index (D). Means ± SD.

Control plots had significantly higher abundance of lichens (all *Cladonia* spp., P-value=0.014), while the N-addition plots had significantly higher Simpson and Shannon diversity values (P-value = 0.0002, P-value = 0.005, respectively; Fig. 2; Table S5a). No significant differences in richness were found between treatments (P-value = 0.633).

PERMANOVA analysis showed significant differences in the lichen communities between treatments (R2 = 0.06, F = 3.0074, *P* value = 0.004, Fig. S5). The species indicator analysis found significant differences for the presence and/or abundance of four species (Table S5b; Fig. S6). *Cladonia crispata (P* value = 0.024), *C. furcata* (*P* value = 0.002) and *C. gracilis* (*P* value = 0.018) were associated with the control plots while *C. fimbriata* (*P* value = 0.002) was associated with N-addition plots. Differences in presence and/or abundance in other *Cladonia* spp. (Table S6), and the lichen crust cover recorded in Southon et al. (2012) were no longer apparent.

### Soil characteristics and leaf chemistry

Soil chemistry analyses revealed significant differences in several variables between treatments; soil pH and Mg were significantly lower in the ammonium sulphate-addition plots (*P* value < 0.01, < 0.05), while CEC was higher (*P* value < 0.01)(Table S7).

The Principal Component Analysis biplot indicates that soil pH, Mg and CEC explain most soil characteristic differences between control and N-addition plots and that Mg strongly correlates with pH (Fig. 3A). The group of soil characteristics are also clustered by control and N-addition, further supporting a difference due to treatment (Fig. 3B). Approximately 47% of the variance is explained in the first (31.3%) and second (16.2%) principal components.

**Fig. 3.**
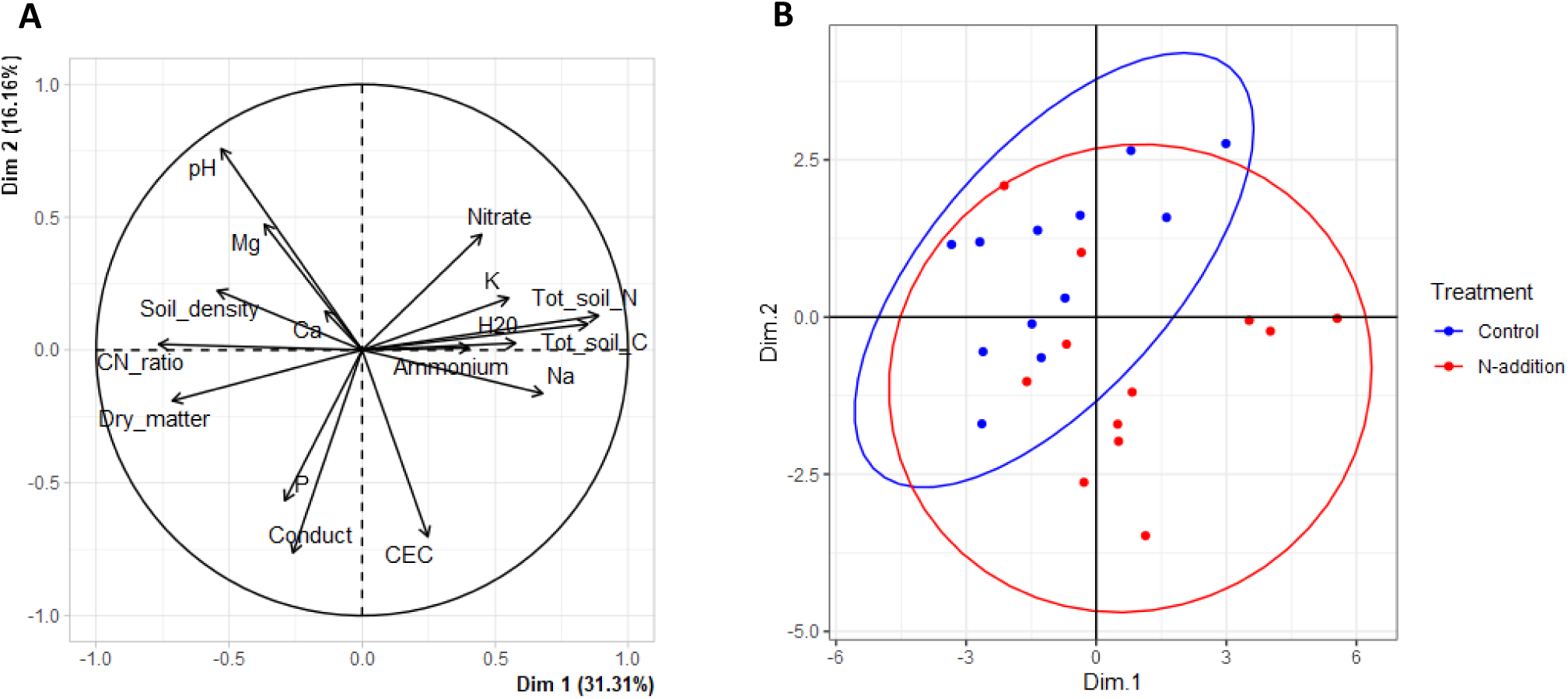
(A) Principal Component Analysis of soil characteristics; (B) Loading plots of soil characteristics based on treatments.

Carbon stock in the control plot samples ranged from 13 to 40 C t/ha while the ammonium sulphate-addition plots ranged from 16 to 48 C t/ha but this was not significantly different. Foliar P, N and C were not significantly different between the treatment plots (Table S8).

### Root ErM fungal colonisation, richness and diversity

#### ErM fungal colonisation

All *Calluna vulgaris* roots analysed showed some ErM colonisation. The analysis of treatment impact on proportion of individual roots colonised, by block and plot, was not significant (Fig. 4A). However, when considering the proportion of overall root cells colonized in all plots combined, the N-addition plots had significantly lower colonisation than control plots (Fig. 4B, Table S9).

**Fig. 4.**
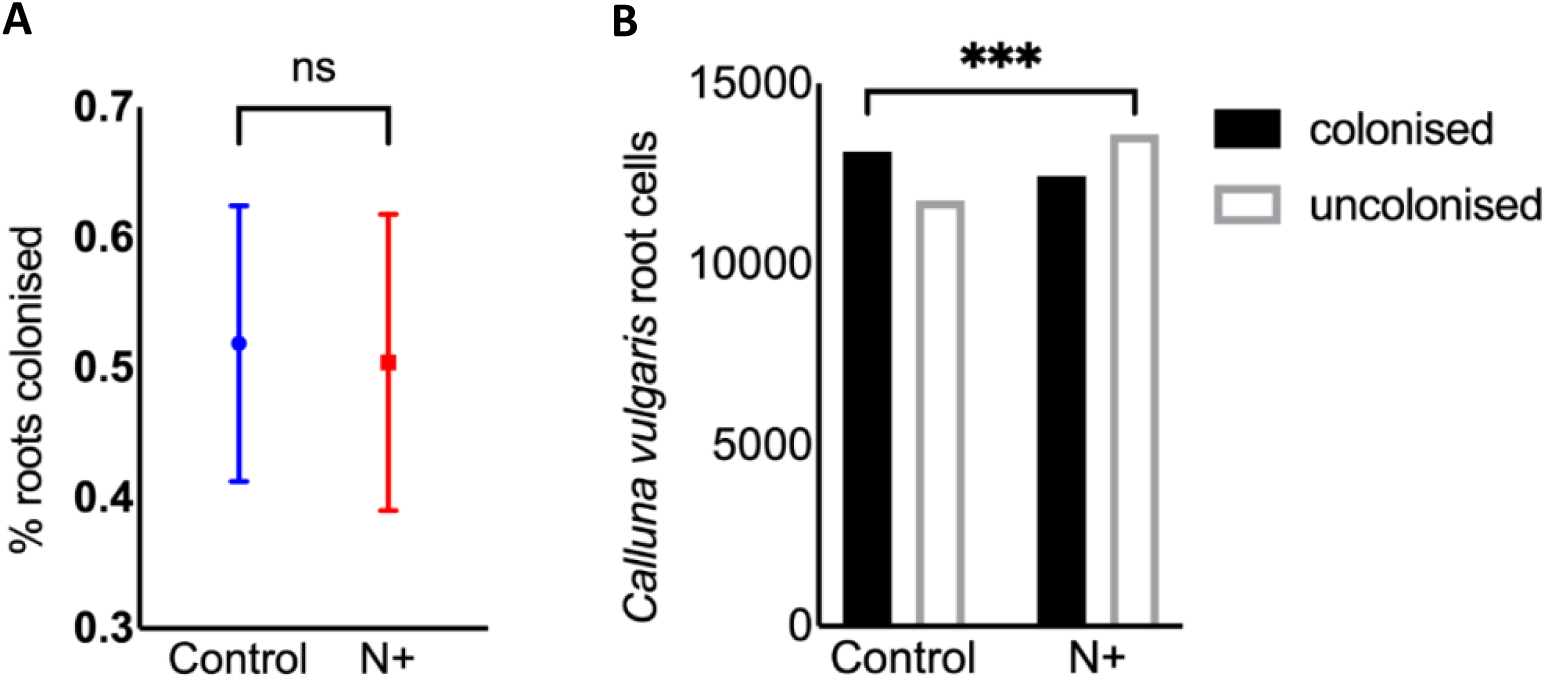
Root colonisation by treatment analysed (A) as mean percentage of individual roots colonized (*p* > 0.05); and (B) Chi-square of all *Calluna vulgaris* cells evaluated (*X*^2^(1, *N* = 50,918) = 11.2, *p* < .0001). Significant *P* value differences are shown as: ***, < 0.001, **, < 0.01; * < 0.05, ns = not significant. Means ± SD.

#### Root fungal richness and diversity

A total of 352 fungal full ITS sequences out of the 368 sequences identified passed the chimera detection and quality control steps. These were clustered into 218 ASVs yielding 23 unique SHs. From these, 56 were assigned as putative ErM ASVs matching to seven unique ErM SHs (Table S10). Three of the seven unique ErM SHs were present in the roots and soil (this study) and sequences from Van Geel et al. (2020) analysed for comparison in this study.

Taxonomic classifications for the following order and families were assigned as: (1) ErM - Chaetothyriales, Helotiales, *Hyaloscypha*/*Pezoloma*, *Hymenoscyphus, Meliniomyces; Oidiodendron*, Sebacinales and Serendipitaceae (Sebacinaceae isolates however have not been resynthesized in culture, Allen et al. (2003)); and (2) ericoid associates - Mycenaceae and Hymenochaetales. Although their mycorrhizal status remains uncertain, some fungi are recognized as frequent associates of Ericaceae roots and are therefore differentiated herein as ‘ericoid associates’.

The ErM fungi represented ∼ 19% and 29% of sequences for the control and N-addition plots, respectively. The different taxonomic breakdown of putative ErM and non-ErM fungi can be seen in Fig. 5A, C.

**Fig. 5.**
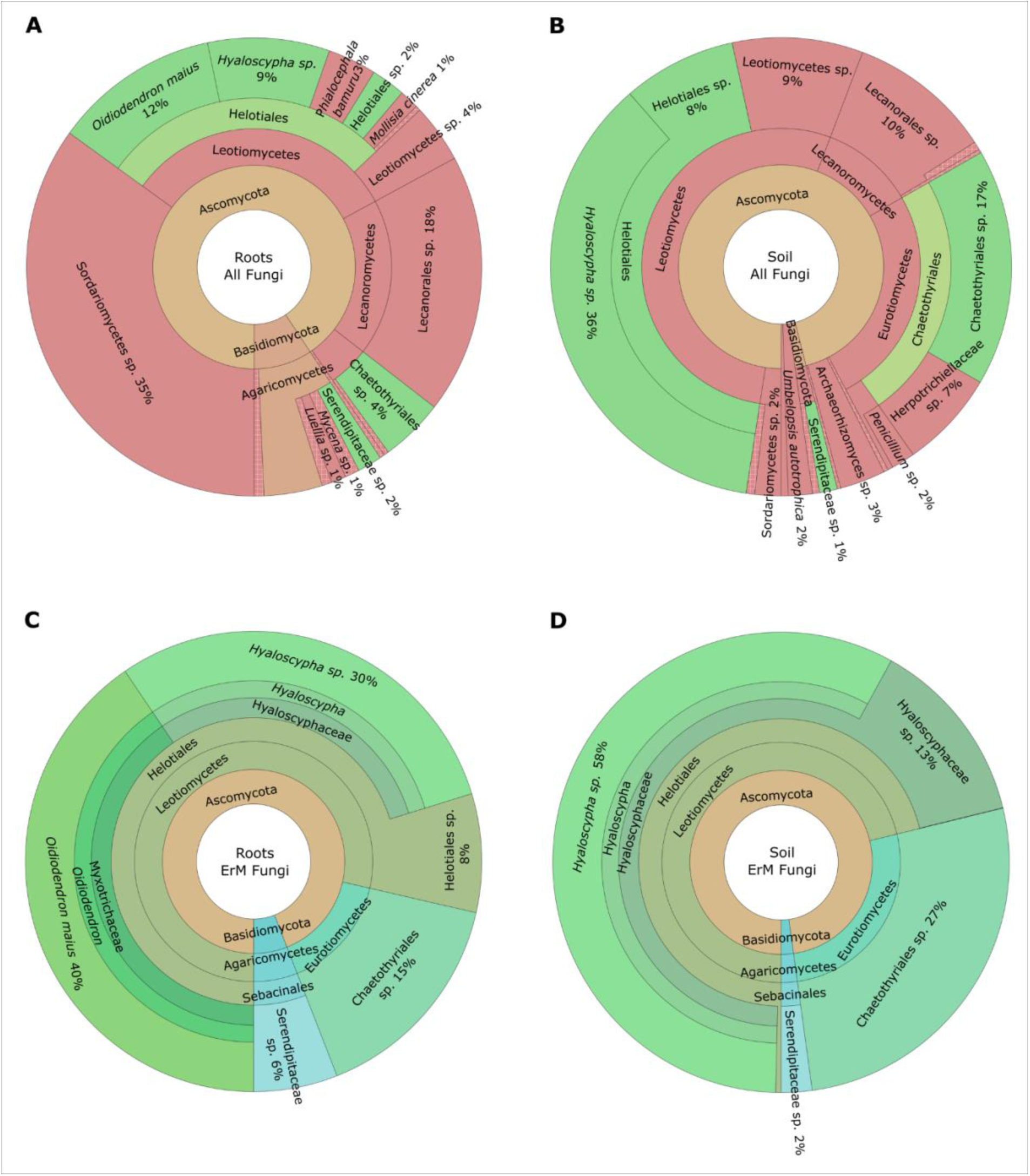
Krona diagrams showing all (A-B) and ErM-only (C-D) fungal taxa identified within *Calluna vulgaris* roots (A-C) and in soil (B-D). The ErM taxa are highlighted in green in charts A and B. Percentages represent the abundance of each taxon in relation to the total of all (A) and ErM only (C) fungal community in roots, and to the dominant soil community taxa (>10,000 reads) in soil (B and D).

Among the first five dominant SHs were the ericoid mycorrhizal genera *Hyaloscypha*/*Pezoloma* (SH1156094.08FU, 22 ASVs) and *Oidiodendron* (SH1185398.08FU, 11 ASVs), which represented 6.4%/7.7%, and 7%/12.1% of the relative abundance in control/N-addition plots, respectively (Table S9). Our SH re-analysis with the 239 putative ErM OTUs previously reported in *C. vulgaris* across 21 European heathlands (van Geel et al. 2020) yielded 82 unique ErM SHs (Table S11).

There were no significant differences in community composition between treatments when comparing all root fungi or the ErM only (Fig. 6A, B).

**Fig. 6.**
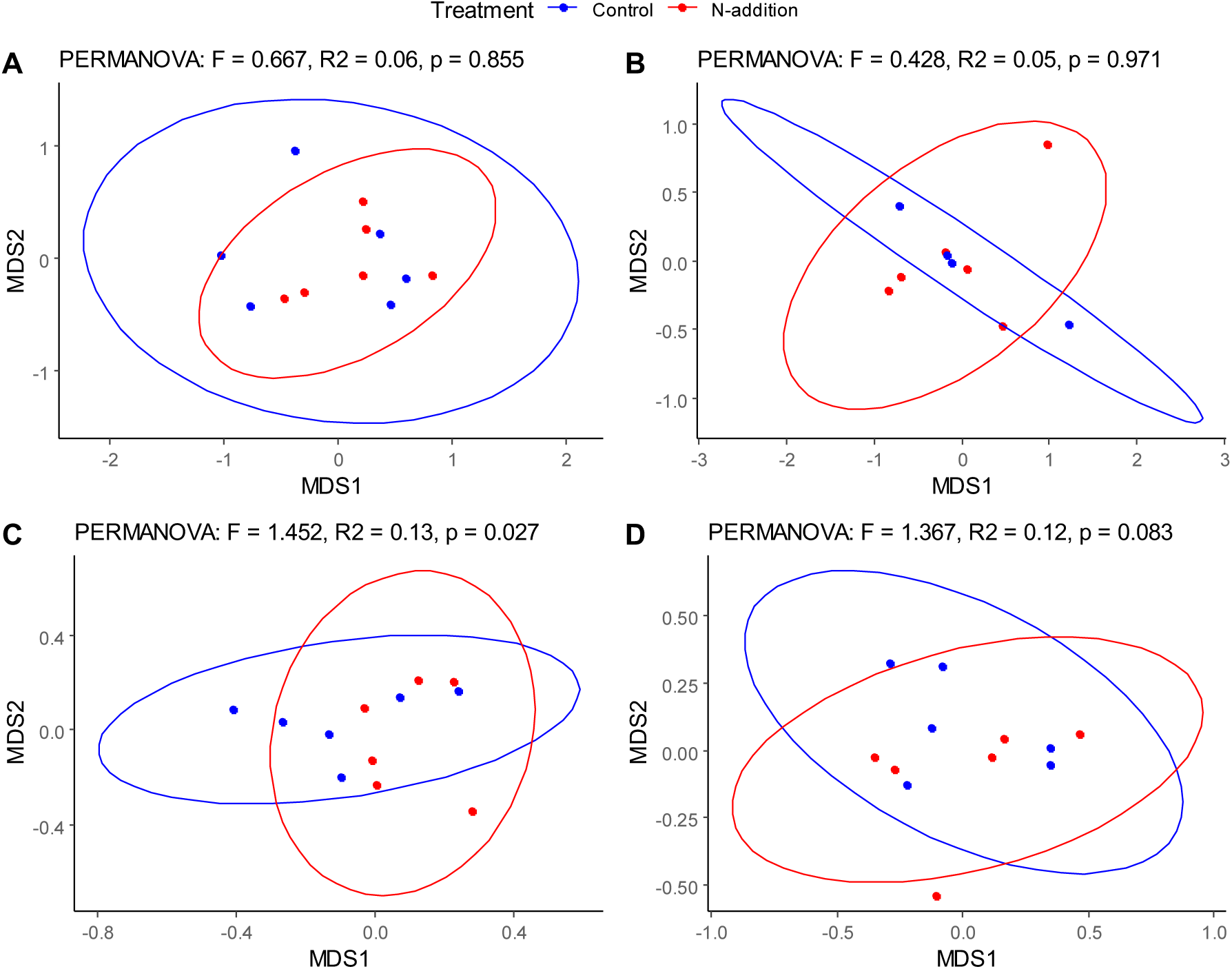
Non-metric multidimensional scaling (NMDS) and PERMANOVA analyses of root (A,B) and soil (C,D) fungal community differences between control and nitrogen (N)-addition treatments. A) All root fungi; B) Putative root-derived ericoid mycorrhizal fungi (ErM) only; C) All soil fungi; D) Putative soil ErM fungi only. All NMDS converge with stress < 0.2.

### Soil fungal richness and diversity

There were significant differences in the overall soil fungi community composition between control and N-addition plots (Fig. 6C). Over 53% of all soil fungi reads were putative ErM which clustered into 1,209 OTUs (97% similarity) matching to 570 unique SHs. Of these, 40 were putative ErM SHs (Table S11). When comparing the soil ErM fungi only, the effect of treatment was not significant (Fig. 6D).

#### Ericoid mycorrhizal richness, soil chemistry covariates

The overall regression model combining soil pH and soil C stock predicts soil ErM SH richness (adjusted R^2^ = 0.756, F _(2,9)_ = 18.05, p < 0.001). Individually each variable significantly predicts soil ErM SH richness (b = 291,123, p = 0.002; b = -3101, p = 0.015, respectively).

Pearson r correlation coefficients between ErM root colonization and ErM soil SH with soil and leaf chemical parameters are reported in Fig. S7.

### Diversity of photobionts associated with Cladonia

A total of 214 ITS rDNA photobiont sequences associated with the genus *Cladonia* were generated; all belonged to *Asterochloris italiana* (Fig. S8). Six haplotypes were identified and connected in a single network. One haplotype was restricted to N-addition plots, three to control plots and the other two haplotypes were present in both treatment plots (Fig. S9). The most abundant haplotype was present in 96.2% of all the specimens and equally represented on plots of both treatments. The genetic diversity of photobionts on control plots was higher than in N-addition plots (Table S12); however, the AMOVA showed that most photobiont variation was within subplots (Table S13).

## Discussion

Our results provide novel insights on the long-term effects of ammonium sulphate addition to European lowland heathlands and highlight the hitherto unappreciated potential of mycorrhizal fungi as bioindicators of heathland recovery. We show that ammonium sulphate addition continues to exert an effect on heathland vegetation structure and ecosystem function more than ten years since treatments ceased and 15 years since a wildfire. Marked differences between treated and control plots are apparent in plant biomass and lichen community composition and diversity, soil characteristics and overall soil fungi community composition. As expected, heather height, both in terms of mean height and highest canopy points, continues to show a N effect, indicating that N-addition has a long-term effect on dominant heathland vegetation structure. However, because all the study heather plots were reaching life-cycle maturity when the plots burned, making new seedling recruitment unlikely (Schellenberg & Bergmeier 2022) it is possible that plant biomass responses and heather regeneration may have been different if the heather stands had burned at an earlier plant life-history stage (Meyer-Grünefeldt et al. 2015). We expected to see comparable growth trends in the stem ring diameters given N additions induce heather canopy growth (Aerts 1989), instead we found that heather stem expansion was insensitive to treatment and that growth regulation was dominated by the terminal stems. Stem thickness and any effects from N addition may have already been reached before the 2006 fire.

In line with previous findings (Terry et al. 2004), our survey also revealed significant differences in lichen community composition more than ten years after ceasing N addition. Thus is likely due to slow growth of lichens (Hale 1959; Karlén & Black 2002) requiring decadal recovery periods.

When considering lichen photobiont diversity, although the number of haplotypes found in the control plots was higher than in the N-addition plots, the two most frequent haplotypes were equally present across treatments, all belonging to *Asterochloris italiana*, a photobiont commonly associated with *Cladonia* species in Europe (Pino-Bodas & Stenroos 2021) and able to tolerate pH values of around 4 (Škvorová al. 2022). Recent studies identified differences in soil chemistry, including pH, N and C content, as major factors driving photobiont switching (Škvorová et al. 2022). However, we found no clear evidence of photobiont switching between our treatment groups, perhaps indicating that treatment levels and soil pH differences were insufficient to elicit such a response.

Turning to bryophyte diversity, N additions are known to promote the invasive moss *Campylopus introflexus* (Sparrius & Kooijman 2013; Caporn et al. 2014; Sérgio et al. 2018), and indeed we recorded significantly higher ground cover of this species in the ammonium sulphate-addition plots. This contrasts with Southon et al.’s (2012) results showing significant differences in species abundance of the moss *Hypnum jutlandicum* and the leafy liverwort *Cephaloziella divaricate* but not of *C. introflexus*. The most likely explanation for this difference is the presence in their study, but not ours, of several other competing pioneer moss species (e.g., *Ceratodon purpureus*, *Pohlia nutans, Polytrichum juniperinum*). Post-fire bryophyte succession on a typical NVC H2 heathland predominantly includes a mosaic of *C. introflexus, C. pyriformis, Ceratodon purpureus, Marchantia polymorpha* ssp. *ruderalis, Pohlia nutans* and *Polytrichum* spp., progressing to dominant *Campylopus* spp. and *Hypnum jutlundicum* by year 20 (Duckett et al. 2008; Burch 2009; Pressel et al. 2021). This is consistent with our observations that only *C. introflexus* and *H. jutlundicum* were present in the experimental plots. Southon et al. (2012) noted that *Cephaloziella divaricata* was significantly more abundant in the control plots. While we did observe *C. divaricata* in many of the plots, we excluded this species from our analyses because it is miniscule and only detectable by close examination with a hand lens, making it impractical in the context of this study. Given the potential of leafy liverworts such as *Cephaloziella* as sources of mycorrhizal inocula to improve the re-establishment of vascular plants following disturbance (Kowal et al. 2016), further studies are needed to determine the long-term effects of N-deposition on this genus.

Surprisingly, observed aboveground differences in bryophyte abundance did not correspond to differences in current soil nitrate or ammonium concentrations. It is likely that given the sandy nature of heathland soils, ammonium and nitrate have been leached, but lingering effects on soil pH remained, since we found significant differences in soil pH values between our control and ammonium sulphate plots, in line with previous reports (Kadyampakeni et al. 2018). The significant differences documented in this study in soil fungal community composition between treatments, but equivocally in mycorrhizal fungal colonisation of *Calluna* roots, further highlight that more experiments are needed to guide policy makers on which components and levels of pollution can lead to fundamental changes in mycorrhizas and C stocks. Differences in pH were the most important predictor of ErM abundance in soil. While N deposition has been linked to lower soil pH, further direct experimental measurements are needed to test this hypothesis in heathland. Southon et al. (2012) reported increased microbial biomass in N-addition plots possibly reflecting greater presence of bacteria and non-mycorrhizal fungi. In this light, our regression results predicting higher ErM abundance when the C stock is lower, are intriguing as they may indicate subtle feedback cycles not captured in our analysis, possibly related to the leaching properties of sandy dry heathland soil. A more plausible explanation is the inherent limitations imposed by the mature age of the canopy which reduces C sequestration potential (Field et al. 2017). Thus, the age of the heather host and relative contribution to soil C stock may yet be another important covariate not measurable in the current study design. Variation in soil C stock across plots may also be explained by uneven distribution of sandy soil and soil moisture (Li et al. 2022), which we found strongly and positively correlated with C stock.

Differences in soil pH can also directly lead to changes in community composition of bryophytes and terricolous lichens (Rai & Gupta 2022). However, all terricolous lichens recorded in our survey belonged to the generally heliophilous genus *Cladonia* (Ahti 2000). Indicating that the most important factor influencing differences in *Cladonia* species richness between the control and ammonium sulphate-addition plots was the density of the heather canopy, as measured by Southon. The indicator species of the ammonium sulphate-addition plots was *C. fimbriata*, which was mainly present at the base of heather near ground level. This species is common in a wide diversity of forests (Burgaz et al. 2020) and has been associated with disturbed sites in the UK (Pino-Bodas et al. 2021). Indicator species in the control plots included *C. crispata*, a characteristic species of mature heathlands (Coppins & Shimwell 1971; Sanderson 2017) and *C. furcata*, a species growing in a wide range of habitats, also frequent in heathlands.

Our molecular analyses of heather root-derived fungal DNA detected 40 unique SHs, of which seven belonged to putative ErM fungi, compared to 82 SHs in heather across 21 dry European heathlands including nine in England (van Geel et al. 2020). Van Geel et al.’s sampling effort i.e., 10 roots from three plants (for each of the three 1 x 1 m plots per site) was similar to that in this study. Six out of the seven ErM fungi found in the roots and 24 out of the 40 ErM fungi detected in soil at Thursley were also identified in their study. Thus, the diversity of ErM hosted by heather at Thursley yielded only one new SH not previously detected at the regional scale.

Our results showing a weak positive relationship between ErM colonisation and P concentration in heather leaves reflect slower growth rates in the aging experimental heather stands alongside less heather regeneration (Calvo-Fernández, 2018). We also found that soil ammonium was negatively correlated to soil putative ErM taxa, but are unable to infer whether this is a consequence of the long-term ammonium sulphate addition experiment since we found no measurable link between soil N and ErM colonisation of *Calluna* roots. ErM colonisation may have recovered to pre-treatment colonisation levels or perhaps was never affected by the treatment. Nevertheless, our observed differences in soil pH and soil fungal community composition between treatments however indicate that treatment effect is likely in this case and is consistent with Van Geel et al. (2020). It is worth noting that N deposition has been previously shown to increase P acquisition (and K) in heather (Rowe et al. 2008), thus it is possible that the weak positive correlation between ErM colonisation and foliar P is a lingering effect of past N additions and that our sample size was simply too small to detect a significant difference between treatments. That soil C stocks in both the Scottish and this study (at least in the top 15 cm of soil) showed no significant differences between N treated and control plots is probably attributable to considerable leaching in this habitat, as discussed before. This, as well as soil pH should be tested in further experiments as explanatory variables for ErM colonisation, diversity and functionality. It is also possible that cumulative historical background N deposition had already impacted the heathland belowground to a level whereby it is difficult to detect strong soil carbon or mycorrhizal effects, despite adding massive amounts of ammonium sulphate and then stopping. Thus, a belowground tipping point may have been crossed long ago whereby the control is largely indistinguishable from treatment.

Despite heathlands harbouring significant biomass belowground, there is a scarcity of experiments testing the effects of atmospheric pollution on these habitats belowground. The fact that differences in vegetation biomass, lichen composition, soil chemistry and mycorrhizal fungal community recorded here can be attributed to the historical ammonium sulphate applications is notable given that mean annual precipitation and temperatures at Thursley have fluctuated dramatically since 2009 (Fig. S1). This suggests that heathlands may be an ecosystem resilient to climate change but more sensitive to deposition of nutrients. Our study thus provides a baseline for monitoring the above and belowground effects of both climate change and nutrient inputs on lowland heathlands.

Predicting interactions between atmospheric pollution, ErM community composition and C sequestration is particularly complex as ErM can be free-living saprotrophs, and their function is context-dependent and strongly related to ErM species diversity (Ward et al. 2022). . In summary, the feasibility of developing objective indices based on ErM abundance and community composition offers a promising new ecological indicator for comparing atmospheric pollution effects and recovery across heathlands in addition to aboveground lichen and vegetation structure and phenology.

## Acknowledgements

Funding for this study came from Kew Science Pilot Grant scheme, the Joint Nature Conservation Committee and the Ecological Continuity Trust. R. P-B. was funded by the program ‘Atracción de Talento Investigador’, Comunidad de Madrid (Spain), 2020-T1/BIO-20503. We thank Emma Green and Georgina Southon for providing their time and raw data from previous long-term experiment studies at Thursley NNR. We are ever grateful to James Giles, the Thursley NNR site warden and Natural England who continue to support our science investigations by providing access to the site and Bliss Furtado, who helped analyse the heather roots for colonisation levels.

## Data availability statement

Our soil and root-derived fungal sequences have been deposited in National Center for Biotechnology Information (NCBI) (accession numbers: pending) and one sequence per photobiont haplotype has been submitted to GenBank (accession numbers: OQ428652-OQ428657). All other raw data is in supplemental information.

