## Supplemental Information for "Assessing above and belowground recovery from ammonium sulfate addition and wildfire in a lowland heath: mycorrhizal fungi as potential indicators"

1- Royal Botanic Gardens, Kew, England
2- Universidad Rey Juan Carlos, Móstoles, Spain
3- Imperial College London, England
4- Natural History Museum, London, England

**Supporting Tables**

**Table S1** Chronology of previous N-addition experiments at Thursley National Nature Reserve with fire history

| Study Years | N+ rate | Organism variable tested: Results of N+ | Authors |
| --- | --- | --- | --- |
| 1976 | ***FIRE*** | | |
| 1994-1996 | 7±7 and 15±4 kg N ha^−1^ yr^−1^) | 1. *Calluna vulgaris* (heather) canopy: increased gaps; shoot water loss rate faster 2. heather beetles: adult weight increased leading to insect damage | Power et al. 1998. *New Phytol* 138. |
| 1989-1996 | 0 or up to 15±4 kg N ha^−1^ yr^−1^ | Recovery after 7 years:   1. heather growth: persistent differences in growth and phenology; suppression of lichen abundance; stimulation of mosses 2. soil pH: recovered 3. soil microbial community size/activity | Power et al. 2006. *Glob Change Biol* 12. |
| 1998-  ***2006 FIRE***  2010 | 0 or 30 kg N ha^−1^ yr^−1^ | Half the original plots (4 of 8 blocks) but double the N-addition:  1. drought injury: exacerbated  2. heather growth: persistent differences in growth and phenology (significantly more bud bursts than the controls over three consecutive years)  3. ongoing suppression of lichens and increased vigour in some bryophytes - the moss *Hypnum jutlandicum* and the liverwort *Cephaloziella divaricata*, but not the majority of mosses (*Campylopus introflexus*, *Ceratodon purpureus*, *Dicranum scoparium*, *Pohlia nutans* and *Polytrichum juniperinum*  4. soil pH reduction; C and N stocks increased (prior and post-fire)  5. soil microbial community size/activity elevated | Jones & Power 2012; Southon et al. 2012. *Glob Change Biol* 18. |

**Table S2a,b,c** Summary of vegetation (a) density measures (b) and lichen species abundance (c) at Thursley National Nature Reserve, NVC H2 habitat (recorded July 2021).

(see excel SI)

**Table S3** Comparison of aboveground mean biomass measures of *Calluna vulgaris*, *Campylopus introflexus* and *Hypnum jutlandicum* by treatment and block using two-way ANOVA contrasted with results measured by Southon et al. (2012). Mean values shown with standard deviation. *P* value significance = *<0.05; **<0.01; ***<0.001; ns, not significant; n/a, not applicable as not tested. N = 48.

|  | treatments | |  | factor *P* value | | |
| --- | --- | --- | --- | --- | --- | --- |
| Biomass measures | Control | N-addition | subplot level n | Treatment | | Southon |
| Height (mean cm ± SEM) |  |  |  |  |  |  |
| *C. vulgaris* highest point | 44.9±4.8 | 51.7±3.0 | 24 | ***** | | ****** |
| *C. vulgaris* mean height | 40.5±4.4 | 47.3±2.1 | 24 | ***** | | ** |
| Density of vegetation |  |  |  |  |  |  |
| *C. vulgaris* at 20 cm | 24.1±6.7 | 28.6±4.0 | 40 points | ns | | ** |
| *C. vulgaris* ground level | 26.5±3.6 | 23.3±0.9 | 40 points | ns | | n/a |
| *C.* *introflexus* | 8.9±2.5 | 14±2.8 | 40 points | * | | ns^1^ |
| *H. jutlandicum* | 1.9±1.9 | 6.4±3.2 | 40 points | ns | | *** |
| Stem rings |  |  |  |  |  |  |
| Ring years (widest radius) | ns | ns | 5 stems | ns | | n/a |
| diameter | ns | ns | 5 stems | ns | | n/a |
| growth (mm) first 5 years | ns | ns | 5 stems | ns | | n/a |

**Table S4** Summary of growth ring measures.

(see Excel SI)

**Table S5a** Differences in lichen richness and diversity compared with the previous long-term experiment (LTE, Southon et al. 2012). Values are shown as mean ± standard deviation (n=48). Significant *P* value differences are shown as: ***, < 0.001; **, < 0.01; * < 0.05; ns, not significant.

|  | Treatment | |  | | |
| --- | --- | --- | --- | --- | --- |
|  | Control | N-addition | *P* value | t value | LTE |
| Abundance | 58.83 ± 22.30 | 46.42 ± 22.10 | 0.014* | -2.45 | n/a |
| Richness | 7.83 ± 2.80 | 8.13 ± 2.60 | 0.624 | 0.49 | ns |
| Shannon index | 1.27± 0.45 | 1.55 ± 0.40 | 0.005** | 2.76 | n/a |
| Simpson Index | 2.83 ± 0.99 | 3.98 ± 1.64 | 0.000*** | 0.31 | n/a |

**Table S5b** Results of the indicator species analysis, indicating the lichen species associated with the different treatments, Control and N-addition.

|  |  |  | Treatment | |
| --- | --- | --- | --- | --- |
| Species | Indicator statistic | *P* values | Control | N-addition |
| *Cladonia fimbriata* | 0.446 | **0.002** |  | X |
| *Cladonia crispata* | 0.326 | **0.024** | X |  |
| *Cladonia furcata* | 0.499 | **0.002** | X |  |
| *Cladonia gracilis* | 0.335 | **0.018** | X |  |

**Table S6** Mean abundance of each species per treatment.

|  | Treatment | |
| --- | --- | --- |
| Species | Control | N+ |
| *Cladonia cervicornis* | 0.542 ± 0.977 | 0.750 ± 1.511 |
| *Cladonia chlorophaea* | 0.167 ± 0.482 | 0.458 ± 1.021 |
| *Cladonia ciliata* | 0.083 ± 0.408 | 0.120 ± 0.4482 |
| *Cladonia coniocraea* | 0.000 | 0.042 ± 0.204 |
| *Cladonia crispata* | 10.708 ± 13.795 | 3.583 ± 5.641 |
| *Cladonia cryptochlorophaea* | 0.042 ± 0.204 | 0.042 ± 0.204 |
| *Cladonia diversa* | 1.458 ± 1.769 | 2.208 ± 3.050 |
| *Cladonia fimbriata* | 0.875 ± 1.154 | 6.250 ± 7.697 |
| *Cladonia floerkeana* | 1.083 ± 1.586 | 2.625 ± 4.576 |
| *Cladonia furcata* | 27.917 ± 14.108 | 14.667 ± 8.820 |
| *Cladonia glauca* | 0.958 ± 1.706 | 0.750 ± 1.294 |
| *Cladonia gracilis* | 0.750 ± 1.225 | 0.125 ± 0.338 |
| *Cladonia macilenta* | 0.833 ± 2.099 | 1.125 ± 1.650 |
| *Cladonia merochlorophaea* | 0.042 ± 0.204 | 0.000 |
| *Cladonia novochlorophaea* | 0.167 ± 0.482 | 0.458 ± 2.245 |
| *Cladonia polydacdactyla* | 0.000 | 0.167 ± 0.482 |
| *Cladonia portentosa* | 6.292 ± 6.504 | 4.0832 ± 4.138 |
| *Cladonia ramulosa* | 3.125 ± 3.314 | 2.667 ± 3.422 |
| *Cladonia squamosa* | 0.333 ± 1.090 | 0.042 ± 0.204 |
| *Cladonia subulata* | 0.208 ± 0.509 | 0.208 ± 0.509 |
| *Cladonia verticillata* | 3.250 ± 4.406 | 6.042 ± 5.797 |

**Table S7** Comparison of soil characteristics between treatment groups using linear mixed effects models. CEC, cation exchange capacity. Values are shown as mean ± standard deviation (n=24). Significant *P* value differences are shown as: **, < 0.01; * < 0.05.

| Soil characteristics | Control | N-addition | *P* value | t value | df |
| --- | --- | --- | --- | --- | --- |
| pH | 4.70±0.04 | 4.49±0.05 | 0.00** | -4.07 | 10 |
| H_2_0 (mg/l) | 5.53±0.67 | 6.16±0.94 | 0.52 | 8.29 | 10 |
| Total N (% w/w) | 0.04+0.01 | 0.04±0.01 | 0.53 | 0.65 | 10 |
| Total C (% w/w) | 1.29±0.23 | 1.53±0.32 | 0.35 | 0.75 | 10 |
| C:N ratio | 38.24±1.54 | 36.97±1.91 | 0.52 | -0.66 | 8 |
| C stock (t/ha) | 23.44±3.61 | 26.37±5.11 | 0.57 | 0.58 | 10 |
| P (mg/l) | 4.43±0.85 | 5.00±0.61 | 0.37 | 0.94 | 8 |
| K (mg/l) | 9.08±4.82 | 13.53±3.15 | 0.20 | 1.41 | 8 |
| Mg (mg/l) | 11.83±1.07 | 8.33±1.51 | 0.04* | -2.32 | 10 |
| Nitrate (mg/kg) | 0.05±0.00 | 0.06±0.00 | 0.23 | 1.25 | 20 |
| Ammonium (mg/kg) | 0.08±0.03 | 0.14±0.04 | 0.15 | 1.50 | 22 |
| Dry matter (%) | 90.78±1.90 | 88.46±2.40 | 0.27 | -1.18 | 8 |
| Soil density (g/l) | 1217±34 | 1179±32.81 | 0.17 | 1.22 | 8 |
| Na (mg/l) | 5.31±0.39 | 6.09±0.52 | 0.18 | 1.49 | 8 |
| Ca (mg/l) | 31.30±9.31 | 28.36±13.17 | 0.83 | -0.22 | 22 |
| CEC (meq/100g) | 10.46±0.13 | 10.97±0.16 | 0.00** | 3.21 | 20 |

**Table S8** Results of mean foliar N, P and C analyses comparing the different treatments, Control and N-addition, all not significant.

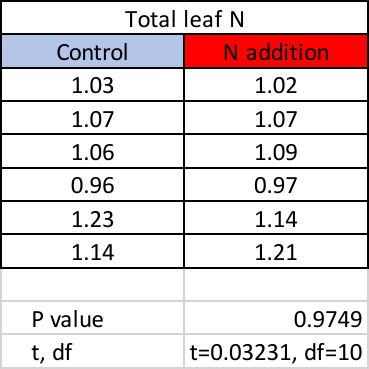

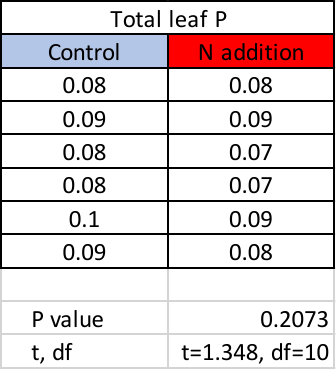

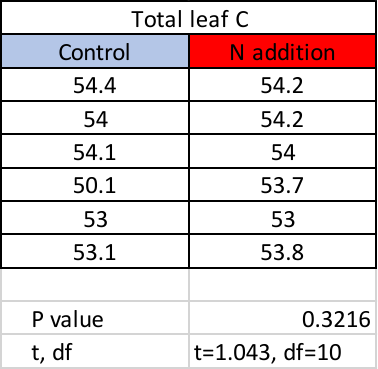

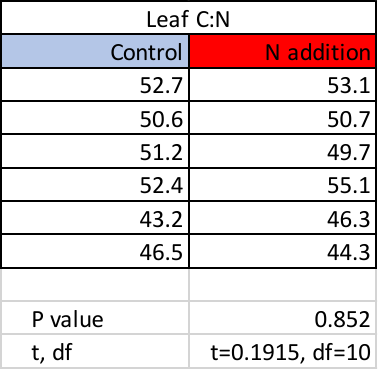

**Table S9** Summary of ErM colonisation count data of all root cells and results of Chi-square analysis (with Yates correction).

| Root cells |  | Control | N-adition | Total |
| --- | --- | --- | --- | --- |
| **Count data** | | | | |
| colonised |  | 13,113 | 12,444 | 25,557 |
| uncolonised |  | 11,773 | 13,588 | 25,361 |
| Total |  | 24,886 | 26,032 | 50,918 |
| **Percentages** | | | | |
| colonised |  | 52.69% | 47.80% |  |
| uncolonised |  | 47.31% | 52.20% |  |
|  | Chi-square degree of freedom | | 121.5,1 |  |
| z |  |  | 11.02 |  |
| P value |  |  | <0.0001**** |  |

**Table S10** Fungal Species Hypotheses (SH) present in *Calluna vulgaris* roots, their taxonomy and putative function, total number of DNA sequences per SH and Amplicon Sequence Variants (ASVs) per SH, by treatment group (Control, N-addition). Relative abundances are calculated as the proportion of sequences in each SH. Green highlights represent ericoid mycorrhiza taxa.

| **Species Hypothesis (SH)** | **SH Taxonomy** | **Putative function** | **#sequences/SH** | | | **#ASVs/SH** | | | **Relative abundance (%)** | | **Presence in soil (this study)** | **Presence in Van Geel et al., 2020** |
| --- | --- | --- | --- | --- | --- | --- | --- | --- | --- | --- | --- | --- |
|  |  |  | **Total** | **Control** | **N-addition** | **Total** | **Control** | **N-addition** | **Control** | **N-addition** |  |  |
| SH1156373.08FU | Sordariomycetes sp. | other | 101 | 63 | 38 | 43 | 28 | 17 | 36.84 | 20.99 | x |  |
| SH1186518.08FU | Lecanorales sp. | lichen | 53 | 24 | 29 | 19 | 10 | 12 | 14.04 | 16.02 |  |  |
| SH1185398.08FU | *Oidiodendron maius* | **ErM** | 34 | 12 | 22 | 11 | 5 | 8 | 7.02 | 12.15 |  | x |
| SH1156094.08FU | *Hyaloscypha finlandica* | **ErM** | 25 | 11 | 14 | 22 | 11 | 12 | 6.43 | 7.73 | **x** | **x** |
| SH1142376.08FU | Rhytismatales sp. | other | 12 | 11 | 1 | 7 | 7 | 1 | 6.43 | 0.55 | x |  |
| SH1186366.08FU | Chaetothyriales sp. | **ErM** | 11 | 6 | 5 | 9 | 6 | 4 | 3.51 | 2.76 |  | x |
| SH1236001.08FU | *Phialocephala bamuru* | other | 10 | 4 | 6 | 7 | 3 | 5 | 2.34 | 3.31 |  | x |
| SH1155585.08FU | Helotiales sp. | **ErM** | 7 | 1 | 6 | 7 | 1 | 6 | 0.58 | 3.31 | **x** | **x** |
| SH2604623.08FU | Lecanoromycetes sp. | lichen | 7 | 2 | 5 | 5 | 2 | 3 | 1.17 | 2.76 |  |  |
| SH2618221.08FU | Agaricomycetes sp. | other | 5 | 5 | 0 | 5 | 5 | 0 | 2.92 | 0.00 |  |  |
| SH1172345.08FU | Serendipitaceae sp. | **ErM** | 4 | 1 | 3 | 4 | 1 | 3 | 0.58 | 1.66 | **x** | **x** |
| SH1156864.08FU | *Mollisia cinerea* | other | 4 | 0 | 4 | 3 | 0 | 3 | 0.00 | 2.21 |  |  |
| SH1172852.08FU | Hymenochaetales sp. | other | 3 | 0 | 3 | 3 | 0 | 3 | 0.00 | 1.66 | x |  |
| SH1170828.08FU | *Mycena romagnesiana* | ericoid_associate | 2 | 2 | 0 | 2 | 2 | 0 | 1.17 | 0.00 | x |  |
| SH1186383.08FU | Chaetothyriales sp. | **ErM** | 2 | 0 | 2 | 2 | 0 | 2 | 0.00 | 1.10 |  |  |
| SH1186231.08FU | *Saitozyma podzolica* | other | 2 | 0 | 2 | 2 | 0 | 2 | 0.00 | 1.10 |  |  |
| SH1170824.08FU | *Mycena zephirus* | ericoid_associate | 1 | 1 | 0 | 1 | 1 | 0 | 0.58 | 0.00 | x |  |
| SH1171997.08FU | *Luellia recondita* | other | 1 | 0 | 1 | 1 | 0 | 1 | 0.00 | 0.55 |  |  |
| SH1155587.08FU | Ascomycota sp. | other | 1 | 0 | 1 | 1 | 0 | 1 | 0.00 | 0.55 | x |  |
| SH1160531.08FU | *Penicillium adametzii* | other | 1 | 0 | 1 | 1 | 0 | 1 | 0.00 | 0.55 | x |  |
| SH1193918.08FU | Serendipitaceae sp. | **ErM** | 1 | 1 | 0 | 1 | 1 | 0 | 0.58 | 0.00 |  | x |
| SH1172222.08FU | Mycosymbioces sp. | other | 1 | 1 | 0 | 1 | 1 | 0 | 0.58 | 0.00 | **x** | **x** |
| SH1171998.08FU | *Luellia* sp. | other | 1 | 0 | 1 | 1 | 0 | 1 | 0.00 | 0.55 | x |  |
| SH_unknown | na | na | 63 | 26 | 37 | 60 | 25 | 35 | 15.20 | 20.44 |  |  |
| **Total** | | | **352** | **171** | **181** | **218** | **109** | **120** |  |  |  |  |
| **ErM (%)** | | | **23.9** | **18.7** | **28.7** | **25.7** | **22.9** | **29.2** |  |  |  |  |
| **non-ErM (%)** | | | **76.1** | **81.3** | **71.3** | **74.3** | **77.1** | **70.8** |  |  |  |  |

**Table S11** Summary of Operational Taxonomic Units (OTUs), Amplicon Sequence Variants (ASVs) and Species Hypotheses (SHs) detected in Thursley root and soil samples compared with our SH compilation of root-derived fungi from 21 European heathlands ‘VG’ (Van Geel et al., 2020).

| **Fungi source** | **OTUs** | | **ASVs** | | **SHs** | |
| --- | --- | --- | --- | --- | --- | --- |
|  | all fungi | ErM | all fungi | ErM | all fungi | ErM |
| Roots (this study, Thursley) | n/a | n/a | 218 | 87 | 23 | 7 |
| Soil (this study, Thursley) | 1209 | 73 | n/a | n/a | 570 | 40 |
| Roots (VG heathlands) | n/a | 239 | n/a | n/a | n/a | 82 |

**Table S12** Summary of genetic variation of photobionts associated with *Cladonia* species on control and N-addition plots.

|  | Control | N-addition |
| --- | --- | --- |
| Number of sequences (N) | 113 | 101 |
| Number of haplotypes (H) | 5 | 3 |
| Haplotype diversity (Hd) | 0.0868 | 0.059 |
| Nucleotide diversity (π) | 0.001 | 0.0003 |

**Table S13** Analysis of molecular variance (AMOVA) within subplot using treatments as grouping factor for ITS rDNA haplotypes of *Asterochloris italiana* associated with *Cladonia*. SS, sum of squared differences of each observation from the mean; F_st_ denotes population differentiation due to genetic structure. ** P-value >0.01; ns = no significant, P-value >0.05.

|  | SS^1^ | Variance components | % Variation | Fixation Index |
| --- | --- | --- | --- | --- |
| Among treatments | 0.024 | -0.001 | -3.03 | F_st_ = 0.128** |
| Among subplots within treatments | 1.147 | 0.005 | 13.23 | F_st_ = 0.102** |
| Within subplots | 6.641 | 0.033 | 89.80 | F_st_ = -0.030 ns |
| Total | 7.812 | 0.037 |  |  |

**Supporting Figures**

**a)**

Thursley Natural Nature Reserve Temperature

|  | J | F | M | A | M | J | J | A | S | O | N | D |
| --- | --- | --- | --- | --- | --- | --- | --- | --- | --- | --- | --- | --- |
| 2009 | 4 | 5 | 8 | 12 | 14 | 17 | 18 | 19 | 16 | 13 | 10 | 5 |
| 2010 | 2 | 4 | 7 | 11 | 13 | 18 | 19 | 17 | 15 | 12 | 6 | 1 |
| 2011 | 4 | 7 | 8 | 14 | 14 | 15 | 17 | 17 | 17 | 14 | 11 | 7 |
| 2012 | 6 | 4 | 10 | 9 | 14 | 15 | 17 | 18 | 15 | 11 | 8 | 6 |
| 2013 | 4 | 3 | 4 | 9 | 12 | 16 | 22 | 19 | 16 | 14 | 7 | 7 |
| 2014 | 7 | 7 | 9 | 12 | 14 | 18 | 21 | 18 | 17 | 14 | 10 | 6 |
| 2015 | 5 | 5 | 8 | 11 | 13 | 17 | 18 | 18 | 15 | 12 | 11 | 10 |
| 2016 | 7 | 6 | 7 | 9 | 14 | 16 | 18 | 19 | 18 | 12 | 7 | 7 |
| 2017 | 5 | 7 | 10 | 10 | 15 | 18 | 19 | 18 | 15 | 14 | 8 | 6 |
| 2018 | 6 | 4 | 6 | 11 | 16 | 18 | 22 | 19 | 16 | 13 | 9 | 8 |
| 2019 | 4 | 8 | 9 | 11 | 13 | 16 | 19 | 19 | 16 | 12 | 8 | 7 |
| 2020 | 6 | 7 | 7 | 12 | 14 | 16 | 18 | 20 | 16 | 11 | 10 | 6 |
| 2021 | 4 | 6 | 8 | 8 | 11 | 17 | 18 | 17 | 17 | 13 | 8 | 7 |
| 2022 | 6 | 8 | 9 | 10 | 14 | 18 |  |  |  |  |  |  |

**b)**

Total annual precipitation Thursley National Nature Reserve

|  | J | F | M | A | M | J | J | A | S | O | N | D |
| --- | --- | --- | --- | --- | --- | --- | --- | --- | --- | --- | --- | --- |
| 2009 | 84.1 | 62.1 | 44.2 | 62.93 | 40.2 | 47.2 | 96.3 | 57.1 | 26.3 | 69.6 | 138.9 | 115.9 |
| 2010 | 80.4 | 103.5 | 48.1 | 35.4 | 27.5 | 33.4 | 34.5 | 100 | 51.7 | 93.4 | 68.8 | 31.1 |
| 2011 | 79 | 64.26 | 29.1 | 14.58 | 46.82 | 102.8 | 96.7 | 65.65 | 32.9 | 41.3 | 38.6 | 74.3 |
| 2012 | 43 | 28.4 | 24.1 | 137 | 85.74 | 194.5 | 99.24 | 54.86 | 52.53 | 119.2 | 130.3 | 110.6 |
| 2013 | 82 | 46.2 | 83.9 | 42.8 | 72.1 | 47.9 | 46.7 | 35.92 | 53.6 | 94.2 | 37.7 | 146.8 |
| 2014 | 99.5 | 150.4 | 40 | 86.1 | 88.3 | 52.1 | 64.4 | 75.63 | 37.74 | 70.3 | 111.13 | 7.62 |
| 2015 | 98.12 | 60 | 31.66 | 24.29 | 69.38 | 35.35 | 82.71 | 72.53 | 24.72 | 66.34 | 80.2 | 87.98 |
| 2016 | 106.39 | 68.31 | 89.34 | 78.2 | 75.94 | 137.21 | 55.33 | 60.2 | 45.7 | 25.81 | 96.26 | 25.36 |
| 2017 | 81.3 | 79.7 | 66.7 | 26.1 | 80.3 | 70.8 | 108.2 | 96.5 | 108.8 | 44.1 | 63.4 | 97.9 |
| 2018 | 73.7 | 43.4 | 130.88 | 58.56 | 112.1 | 10.55 | 97.6 | 64.7 | 28.36 | 37.93 | 99.9 | 104.57 |
| 2019 | 32.5 | 50.3 | 92.6 | 43.8 | 93.1 | 174 | 66.6 | 57.1 | 90.8 | 107.1 | 136.4 | 118.9 |
| 2020 | 80.4 | 95.7 | 33.6 | 41.3 | 10.4 | 84.9 | 29.5 | 87.7 | 40.9 | 122.6 | 76.4 | 78 |
| 2021 | 95.2 | 68.4 | 38.7 | 34.4 | 117.8 | 148.6 | 92.1 | 33.1 | 35.5 | 134.2 | 15.7 | 67.7 |
| 2022 | 22 | 41.9 | 60.2 | 19.3 | 43 | 26.3 |  |  |  |  |  |  |

**Fig. S1**	Mean annual temperature and precipitation (mm), Thursley National Nature Reserve, Surrey (2009-2022, June only).

**a)**

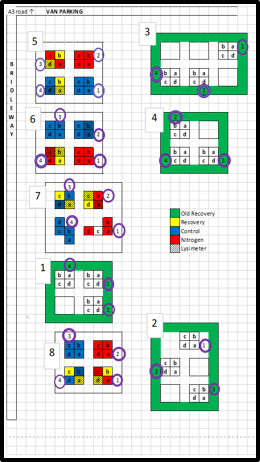

**b)**

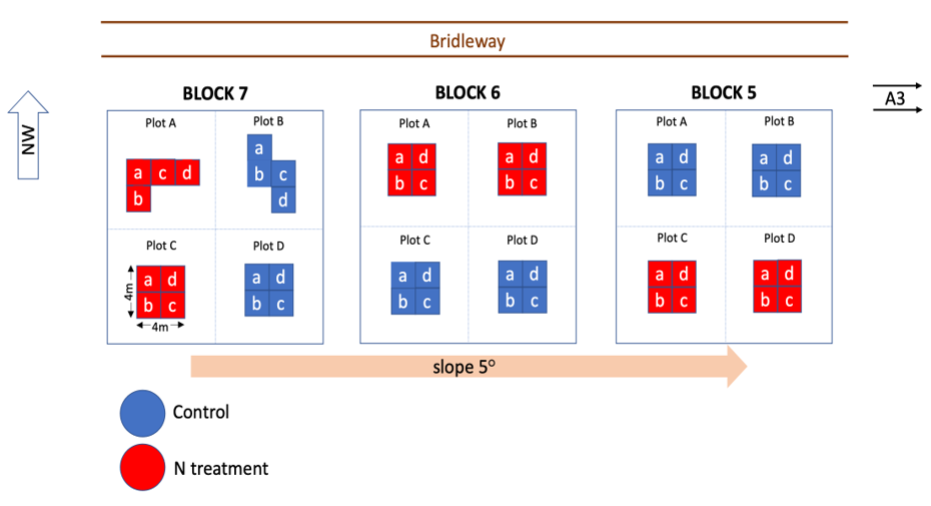

**c)**
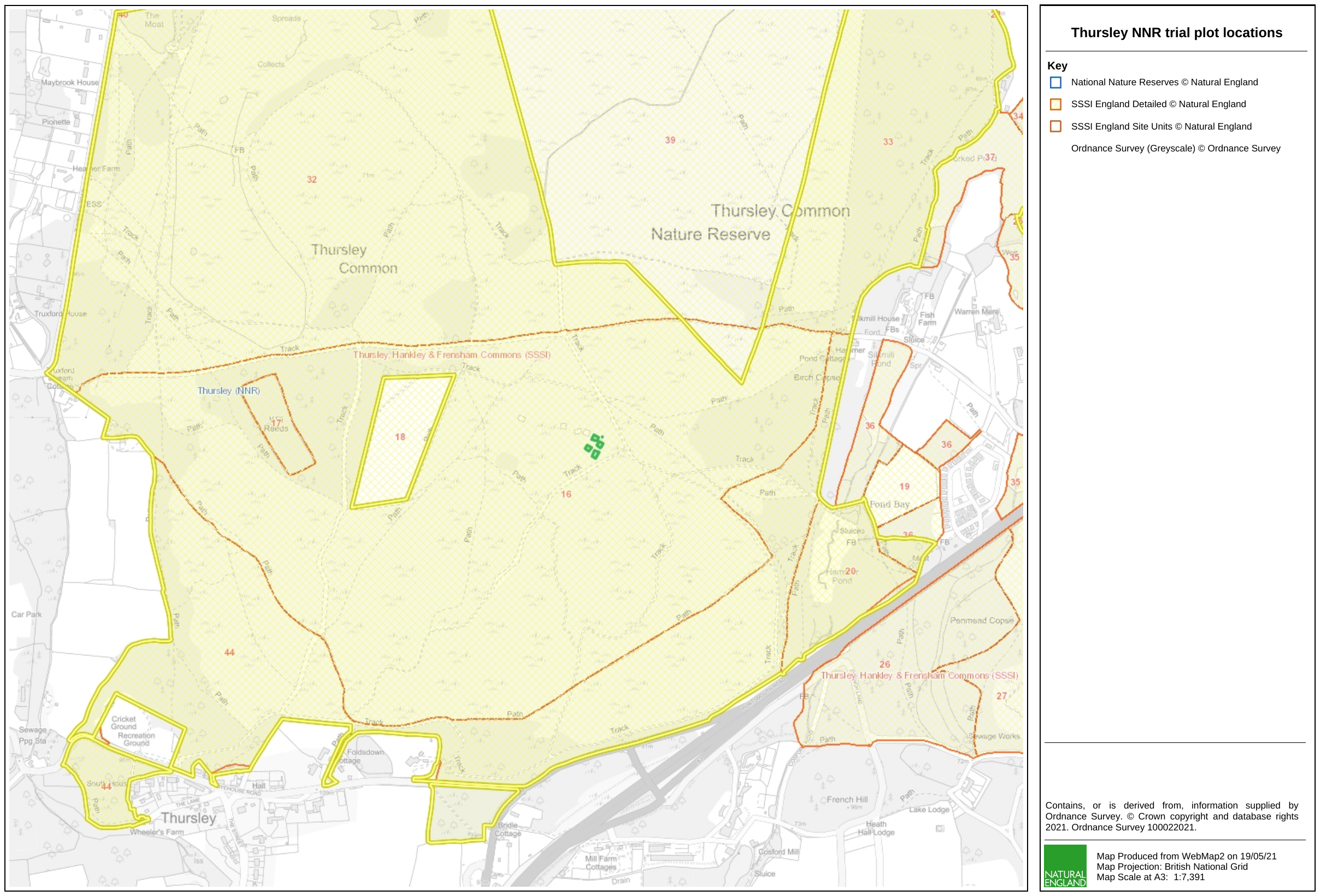

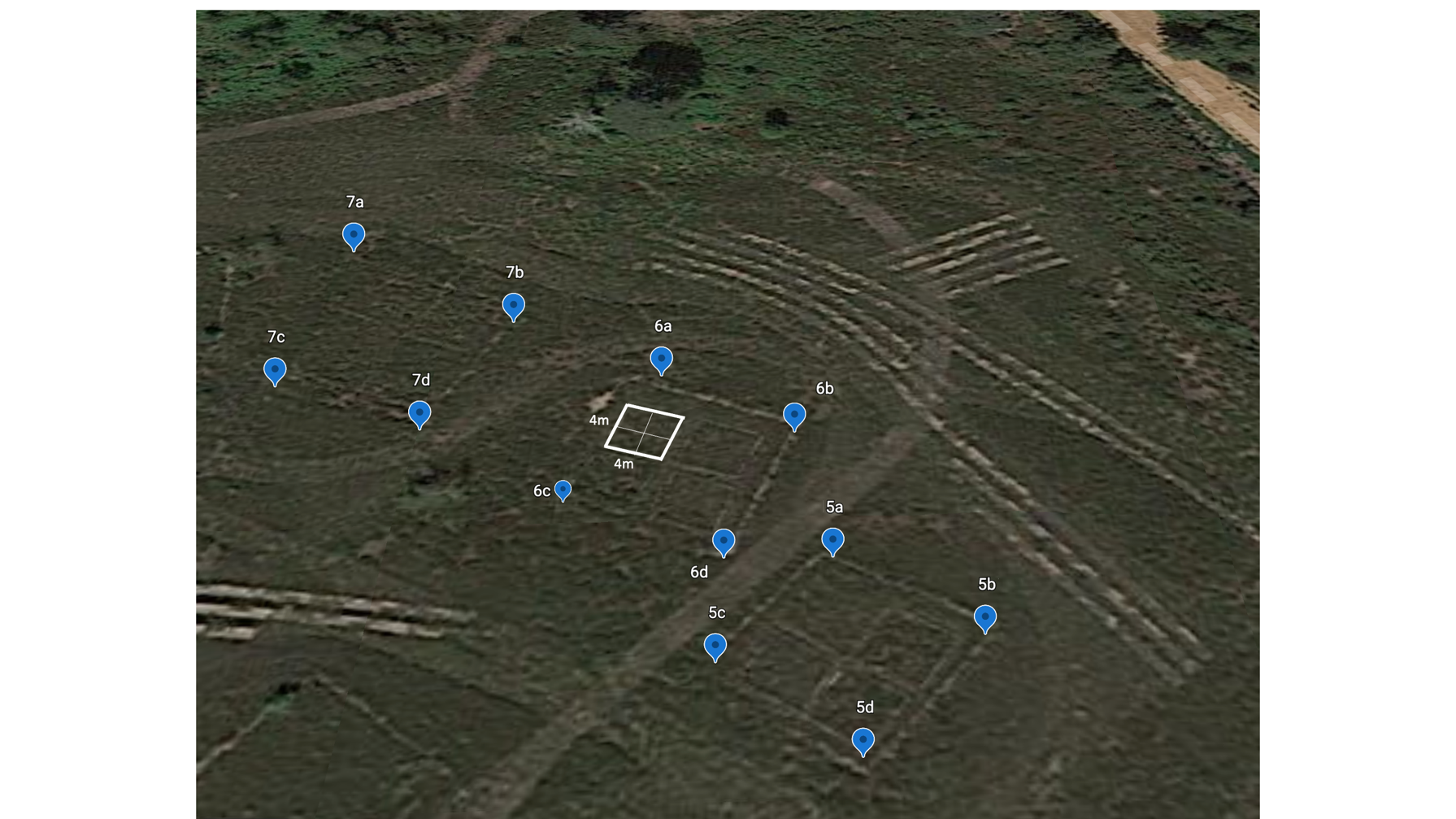

**Fig. S2a, b, c**	(a) Original field experimental block design (Power et al., 1998, modified by Southon et al., 2012). Red Xs indicates blocks removed by Southon. Each block (numbered 1 through 4, in bold) has either six or four 4x4 m plots which are then subdivided into four 2x2 m sub-plots. The management treatments indicated in the key (old recovery and recovery) were not significant. The three blocks tested in this study are circled in red and enlarged in (b). These three blocks each have two ammonium sulphate addition plots and two control. Source: Emma Green notes. (c) Map of Thursley trial plots, circled in red (Source: Natural England) with corresponding aerial view of the plots tested in this study (5,6,7) adapted from Google Earth (downloaded October 2021).

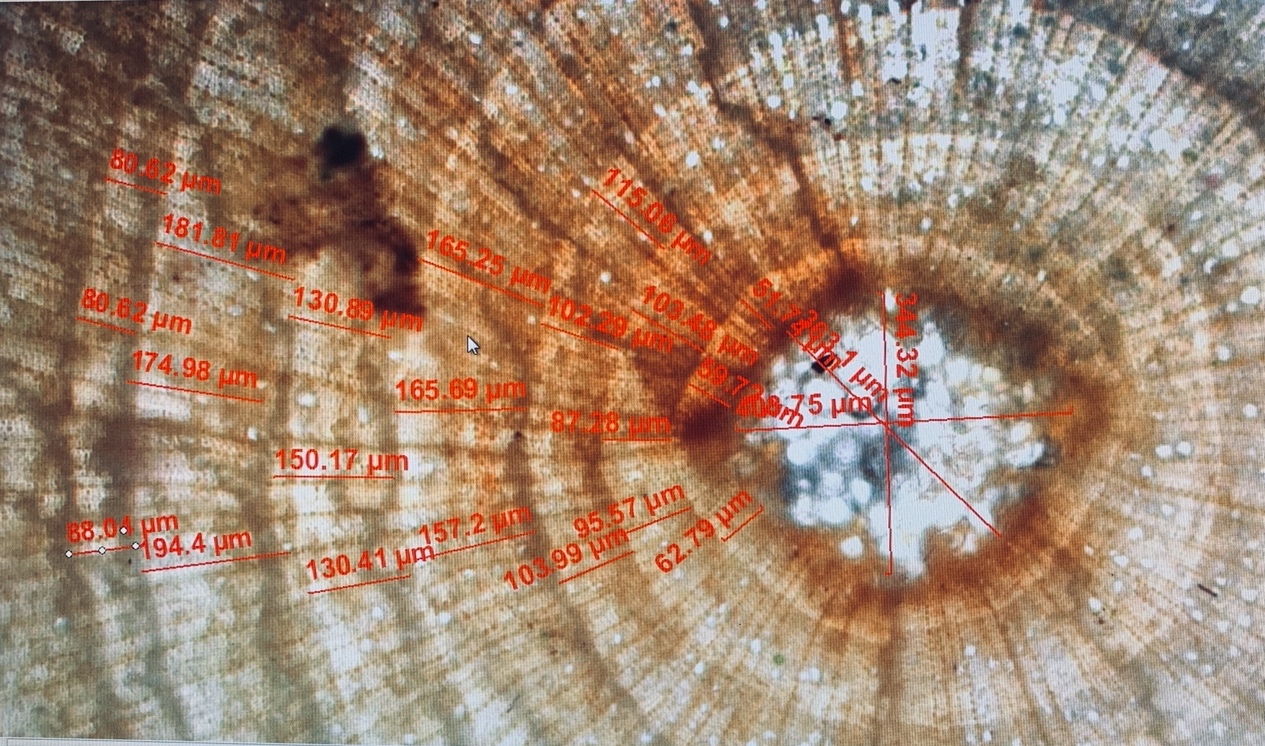

**Fig. S3**	Image showing cross-section of *Calluna vulgaris* stem with three measurements per ring. Pith in the center.

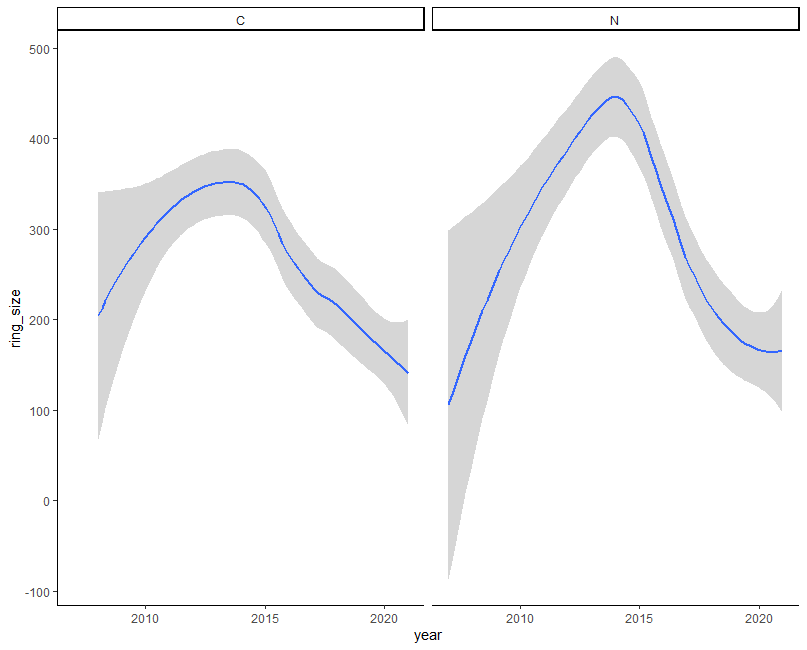

**Fig. S4**	Comparison between control ‘C’ and N-addition ‘N’ treatments of annual ring growth at Thursley post fire of 2006. There is no significant difference in the height of the peak in 2014 between control and N-addition treatments (LMM: t = 0.44, df = 52, p = 0.66).

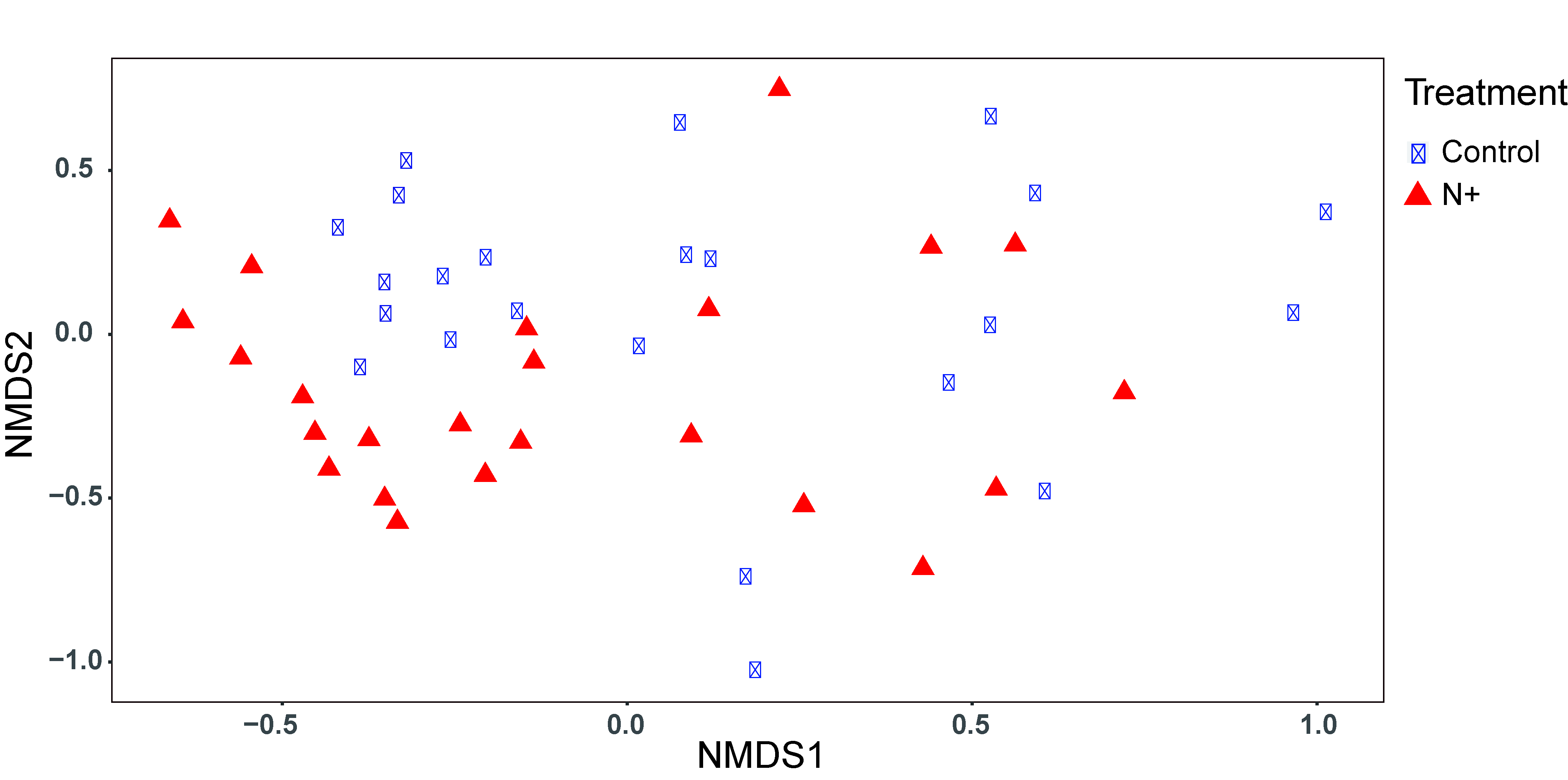

**Fig S5.** Nonmetric multidimensional scaling showing *Cladonia* spp. community based on the Bray-Curtis dissimilarity matrix comparing Control and N-addition (N+) plots. All convergent with stress < 0.2.

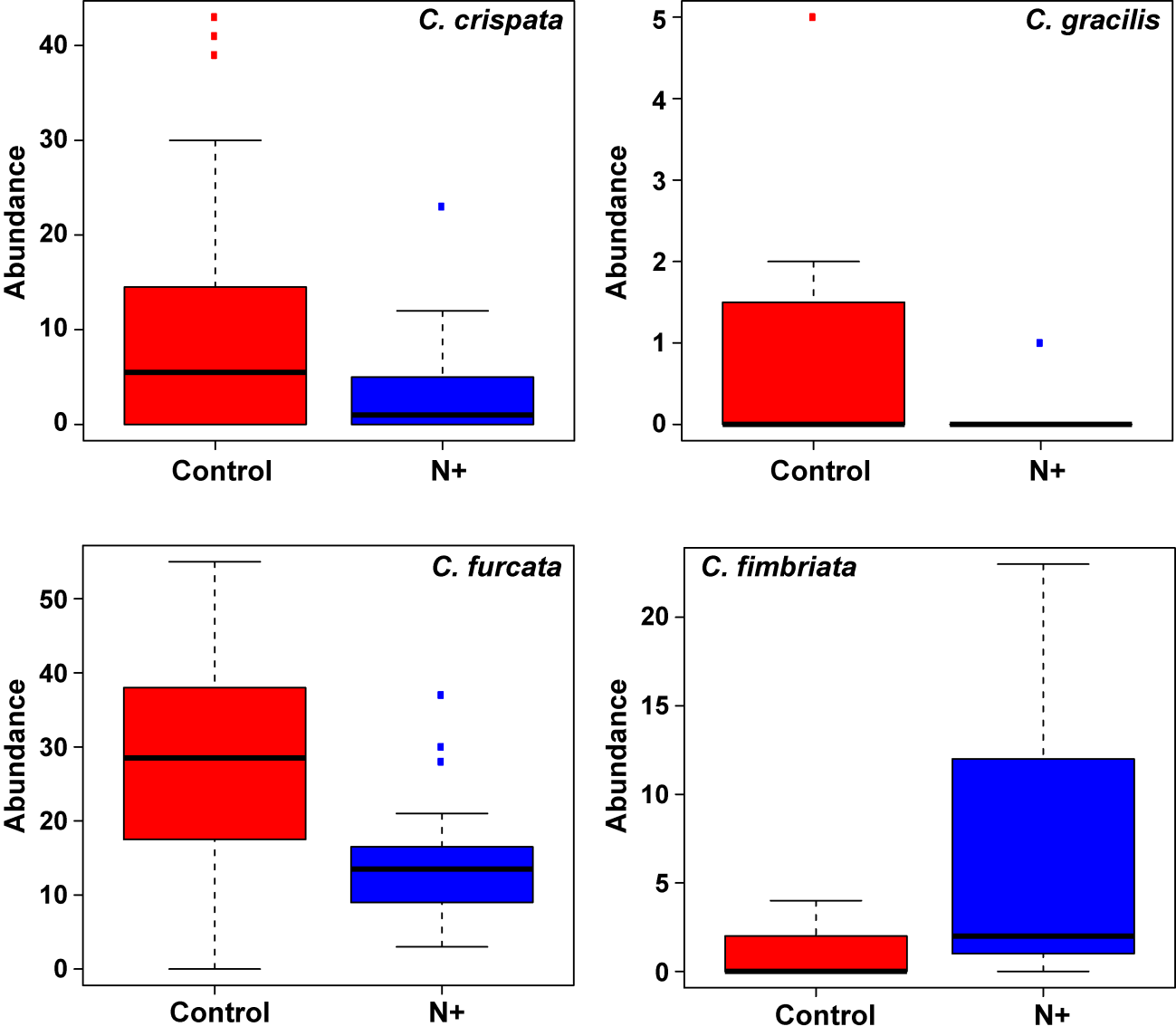

**b)**

**d)**

**c)**

**a)**

**

*

**

*

**Fig S6.** Box plots comparing abundance responses to treatments for four different *Cladonia* spp. a) *C. crispata (P* value = 0.024), b) *C. gracilis* (*P* value = 0.018), c) *C. furcata P* value = 0.002), and d) *C. fimbriata* (*P* value = 0.002); N+, n-addition plots. Means ± SD.

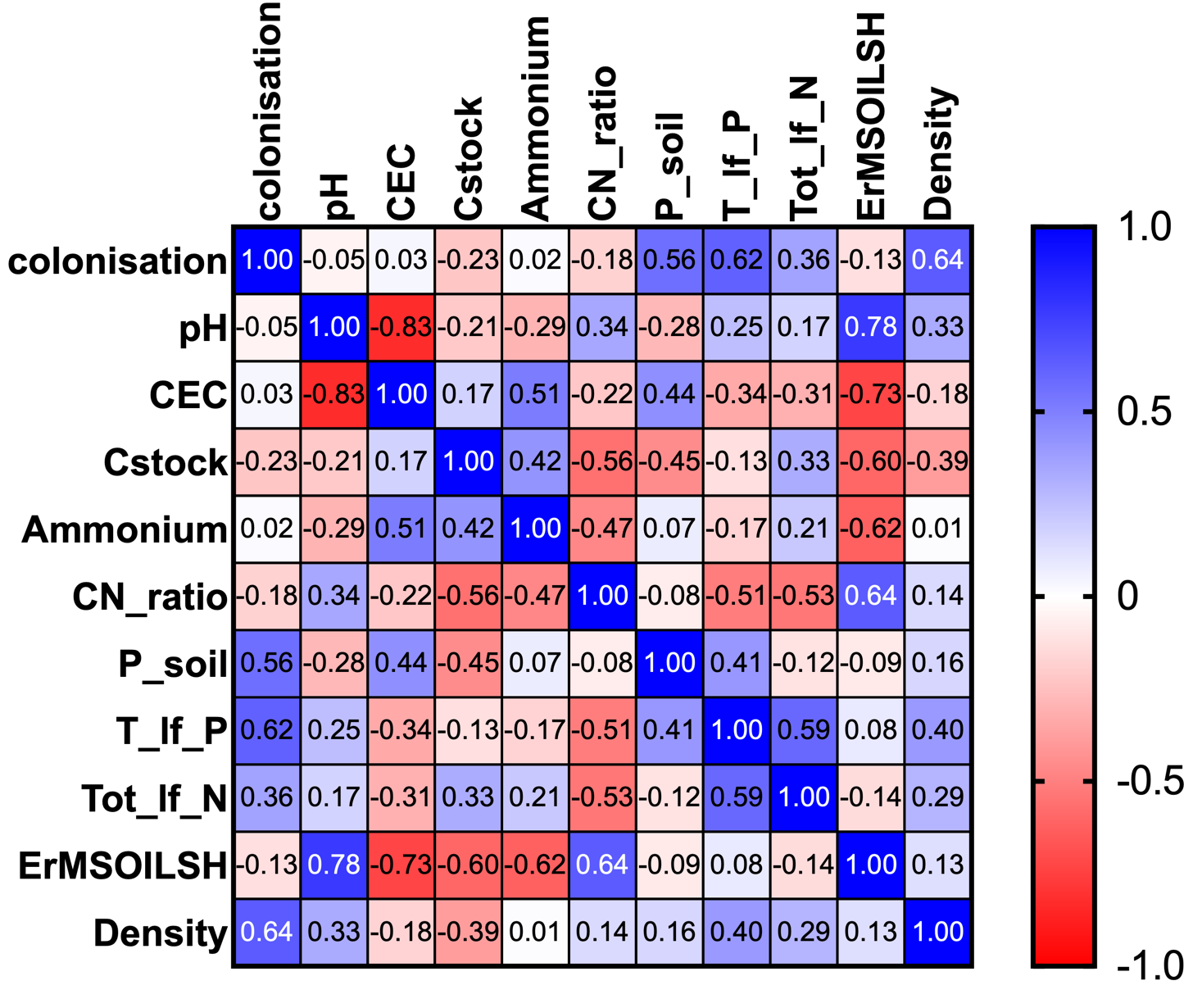

**Fig. S7** Heatmap of Pearson r correlation coefficients between ErM root colonization and ErM soil richness (SH) with soil and leaf chemical parameters.

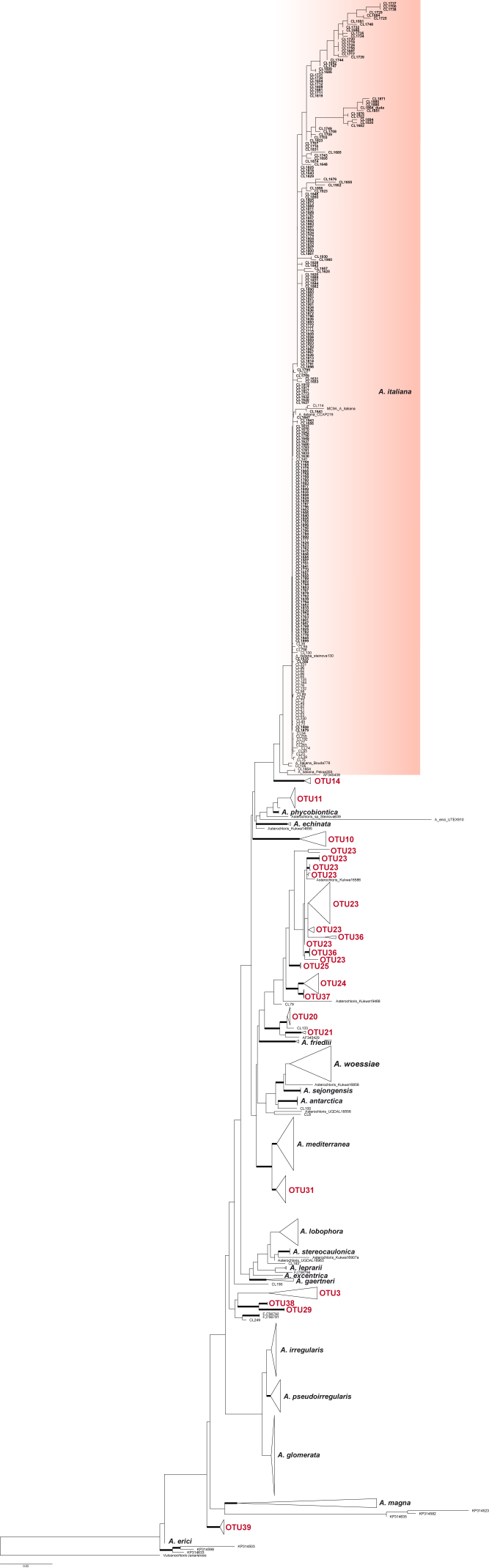

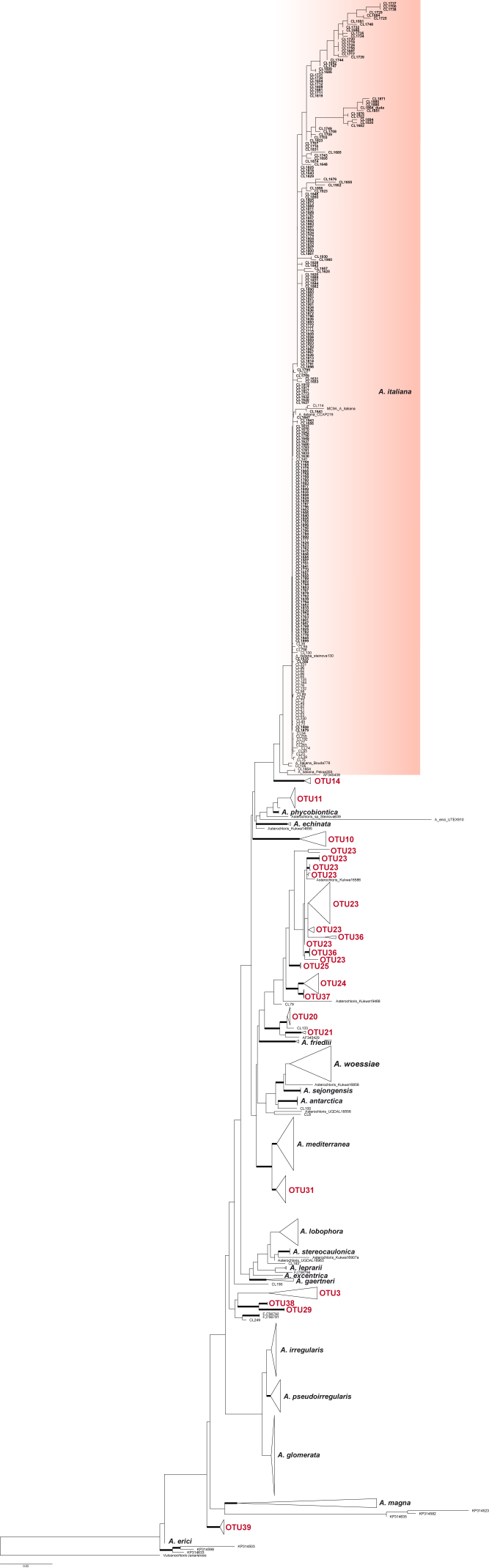

**Fig. S8** Phylogeny of *Asterochloris* based on ITS rDNA. Maximum Likelihood tree of RAxML. Bold branches were supported by ≥ 75% of bootstrap values. The clade including all the sequences of photobionts from the Thursley *Cladonia* spp. is marked in red.

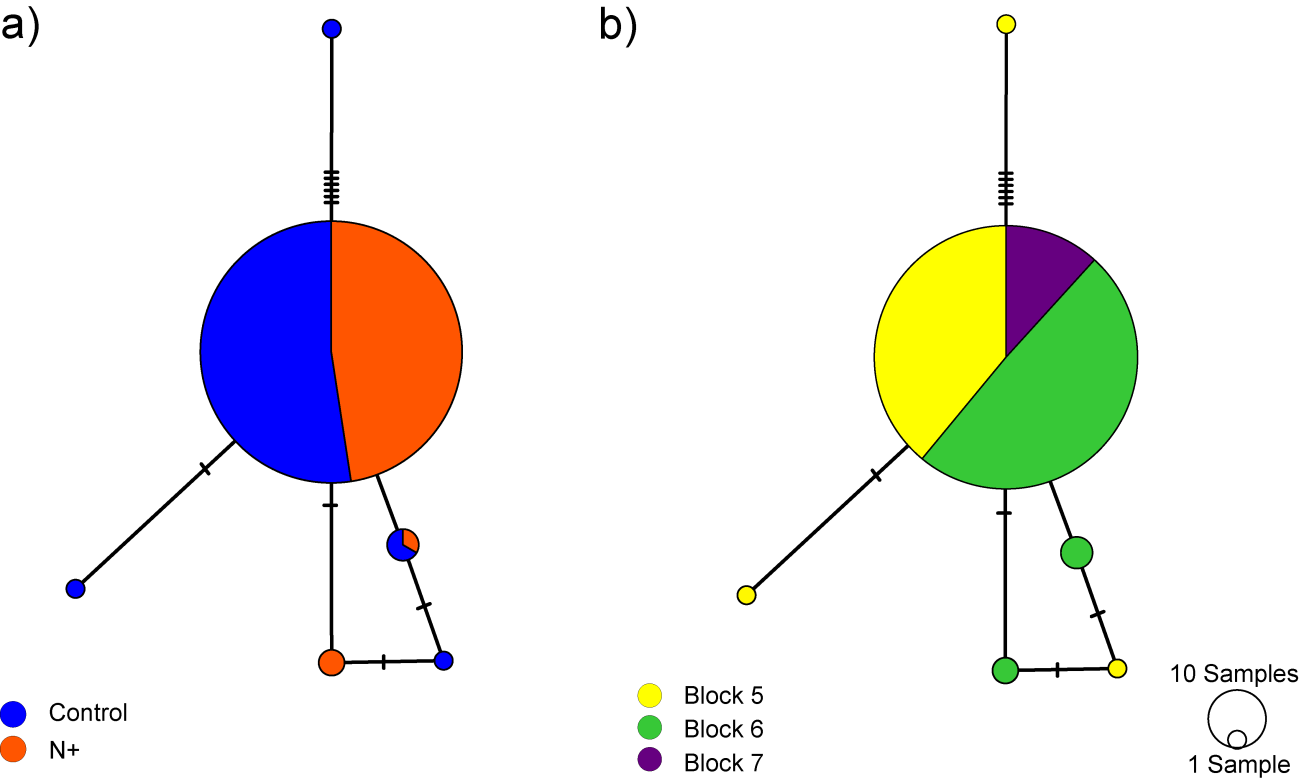

**Fig. S9** Statistical parsimony haplotype network of *Asterochloris* based on ITS. Haplotypes are coloured according to (a) treatment, Control and N+ (nitrogen addition) and (b) blocks 5,6 and 7. The size of the circles is proportional to the frequency of each haplotype. Small circles represent haplotypes not observed in the data.

**Supporting Data S1. Field Experiment: Thursley National Nature Reserve, Surrey, UK (1998-2010)**

In 1989, a long-term experiment was set up at Thursley National Nature Reserve, to mimic and test ammonium sulphate on heathland ecosystem function (Power et al. 2006; Jones & Power 2012; Southon et al. 2012). The results showed that ammonium sulphate addition, applied bi-weekly between 1998 and 2010, resulted in significant changes in aboveground vegetation structure and diversity (Table S1) and belowground nutrient cycling, specifically C and N stocks. In 2006 the study was interrupted by a severe wildfire (Duckett et al. 2008) which affected all experimental plots and completely eliminated the successional heathland structure i.e., pioneer, building, mature and degeneration (Gimingham 1972). Following this event, the effects of fire and N deposition interactions were studied for an additional six years (Southon et al. 2012). Despite the wildfire, which removes most of the available N from *C. vulgaris* heathlands (Pate & Dell 1984; Read 1996), Southon et al. (2012) found continuing effects of the pre-fire N treatments on post-fire vegetation regeneration among pioneer species (Table S1). The dominant heather shrubs no longer had significant differences in growth parameters, likely due to post-disturbance regeneration from older stems (Schellenberg & Bergmeier 2022). The Southon et al. (2012) study also revealed significantly higher soil C in the N-addition plots, which was attributed to increases in the microbial biomass, although it did not examine mycorrhizal colonisation, root-associated fungi or soil fungi community structure.

**Methods S2**. **Soil chemical extraction methods**

Upon returning to the laboratory, visible rocks and roots were removed and soil samples were maintained overnight at 3-4°C. The next day they were delivered in cool boxes to an external laboratory for processing and analyses (NRM Laboratories Services, Bracknell England). Below are the abridged methods employed by NRM Laboratories.

***Determination of soil bulk density***

The total weight of the sample was recorded after drying overnight at < 30°C. This information was used to calculate the soil bulk density on a dry matter basis for the set volume. A scoop of set volume is used to take a sample of the dried and ground soil (after passing through a 2mm sieve). This is then levelled off before the weight is recorded.

Soil Density (g/l) = Weight of soil (g)/Volume of ring (l)

Reference: Soil Science – Methods and Applications. DL Rowell. ISBN 0-582-08784-8.

***Determination of Electrical Conductivity (EC)***

Matrix: Soil samples were air-dried at a temperature <30°C and sieved to pass a 2mm screen, excluding stones and fibrous material and roots.

Soluble salts, other than calcium sulphate, are extracted from soil with saturated calcium sulphate solution. The specific conductivity of the extract at 20°C is recorded as soil conductivity. Results are expressed as uS/cm at 20°C.

Reference: The Analysis of Agricultural Materials, MAFF Reference Book RB427, ISBN 0 11 242762 6.

***Determination of soil pH***

Matrix: Soil samples were air-dried at a temperature <30°C and sieved to pass a 2mm screen, excluding stones and fibrous material and roots.

A suspension of 5 grams of soil with 25 ml of water was shaken on an orbital shaker for 15 min and rested for 45 min prior to pH analysis using a Sentek pH electrode.

Reference method: ISO 10390.

***Determination of total Carbon and Nitrogen***

Matrix: Soil samples were dried and ground to pass through a 0.5mm screen.

Total Carbon and Total Nitrogen are determined by combustion analyser. Results are expressed as percentage in soil.

Reference: AOAC Official Methods of Analysis (1990) Method 949.12. ISO 10694 & 13878.

***Determination of plant available Phosphorus using Melich III* *extractable elements in soil (for soil pH <6.0)***

Matrix: Soil samples were air-dried at a temperature <30°C and sieved to pass a 2mm screen, excluding stones and fibrous material and roots.

The sample is shaken with Mehlich III extracting solution which dissolves the ‘active’ compounds. The extracted compounds are determined by Inductively Coupled Plasma Optical Emission Spectroscopy. This extractant is composed of 0.2M acetic acid; 0.25M ammonium nitrate; 0.015M ammonium fluoride; 0.013M nitric acid; and 0.001M EDTA. Exchangeable K, Ca, Mg, and Na are extracted by the action of ammonium nitrate and nitric acid. Micronutrients are extracted by NH4 and the chelating agent EDTA. Acetic acid and ammonium fluoride are present for the extraction of phosphorus. Results are reported as mg/l in soil on a dry matter basis.

Reference: Mehlich, A. 1984. Mehlich III soil test extractant: A modification of the Mehlich 2 extractant. Commun. Soil Sci. Plant Anal. 15:1409-1416.

***Determination* of *mineral nitrogen in soil (available N)***

Background: The protocol assumes soil available N includes nitrate-N, nitrite-N and ammonium-N that can be extracted by 2M Potassium Chloride.

Matrix: The soil samples were maintained between 2-8°C to restrict nitrogen mineralization which could lead to an overestimation of available N.

The soil sample is chopped and mixed to obtain a homogenous sample, removing stones and roots. A portion is then shaken with 2M KCl to extract the mineral-N fractions and a dry matter determination carried out. Once in solution the Nitrate-N, Nitrite-N and Ammonium-N can be measured colourimetrically. Results are reported as mg/l in soil on a dry matter basis.

References:

Soil Sampling and Methods of Analysis. Canadian Society of Soil Science, Ed Martin R Carter. ISBN 0-87371-861-5.

The Analysis of Agricultural Materials, MAFF Reference Book RB427, ISBN 0 11 242762 6. Reference method ISO 10694 & 13878.

***Determination extractable potassium, calcium, sodium and magnesium in soil***

Matrix: Soil samples were air-dried at a temperature <30°C and sieved to pass a 2mm screen, excluding stones and fibrous material and roots.

The available potassium and magnesium are extracted from the soil by shaking with 1M ammonium nitrate at 20°C for 30 minutes. After filtration, the concentration of potassium and magnesium in the extract is determined by atomic absorption spectrometry. The instrument is calibrated using commercial potassium and magnesium standards traceable to the SI unit.

References:

The Analysis of Agricultural Materials, DEFRA Reference Book RB427 ISBN 0 11 242762 6.

Fertiliser Recommendations for Agricultural and Horticultural Crops, DEFRA Reference Book RB209 9th edition.

***Determination* of *available sulphate***

Matrix: Soil samples were air-dried at a temperature <30°C and sieved to pass a 2mm screen, excluding stones, fibrous material and roots.

Available sulphate is extracted from the soil using a phosphate buffer extracting solution (1:2) and the filtered extract of the sample is analysed by Inductively Coupled Plasma Emission Spectroscopy.

Reference:

Soil Science - Methods and Applications, D L Rowell, 1995, Longman Scientific and Technical, ISBN 0 582 087848.

***Determination* of *available manganese***

Matrix: Soil samples were air-dried at a temperature <30°C and sieved to pass a 2mm screen, excluding stones, fibrous material and roots.

The available element manganese is extracted from the soil at 20°C with DTPA solution (1:2) adjusted to a steady pH of 7.3.

Reference: Developed by Lindsay and Norvell: Soltanpour, P N, and Schwab, A P. 1977. Comm. Soil Sci. Plant Anal. 8:195-207. Handbook on Reference Methods for Soil Analysis - 1992, ISBN 09627606 1 7.

**Methods S3**. **Details of molecular testing and bioinformatics analyses on root-associated and soil fungal communities**

DNA was extracted in duplicate from 250 mg of soil (fresh weight) for each plot, using the DNeasy PowerSoil Pro kit (Qiagen, Hilden, Germany) according to manufacturer's instructions. DNA extracts were quantified using a Quantus^TM^ Fluorometer with QuantiFluor® Dye System for dsDNA (Promega, Madison, Wisconsin, United States) and purity-checked (260/280nm and 260/230nm ratios) via NanoDrop spectrophotometer (Thermo Scientific, Waltham, Massachusetts, United States). DNA extracts were sent to Macrogen (Macrogen Genome Center, Seoul, Republic of Korea) for downstream processing and bioinformatic analysis. PCR amplification of the ITS2 region was performed using the primers ITS1F/ITS86F (White et al., 1990; Turenne et al., 2000), and amplicon libraries were sequenced on an Illumina MiSeq (Illumina, [San Diego, California, United States](https://www.google.com/search?sxsrf=ALiCzsbgjpTrJLUnGWAnyLJMkA4WYbB1pQ:1666106380138&q=San+Diego&stick=H4sIAAAAAAAAAOPgE-LSz9U3MDIvMUxPUeIAsc0Ny4q0tLKTrfTzi9IT8zKrEksy8_NQOFYZqYkphaWJRSWpRcWLWDmDE_MUXDJT0_N3sDLuYmfiYAAAOlk1YlgAAAA&sa=X&ved=2ahUKEwiBu8j3ier6AhWER8AKHZ0jB2wQmxMoAXoECHoQAw)). Reads pre-processing and clustering were carried out by Macrogen Representative sequences from each OTU were assigned to a Species Hypothesis (SH) in house, using the SH matching v2.0.0 tool on PlutoF.

The complete ITS region (ITS1, 5.8S and ITS2) was amplified using the fungal-specific primers ITS1F and ITS4 (White et al., 1990; Gardes and Bruns, 1993). PCR reactions were carried out in a final volume of 10 µl, consisting of 5 µl of JumpStart Taq ReadyMix (Sigma-Aldrich, St. Louis, Missouri, United States), 0.05 µl of each primer (100 µM), 2.4 µl of PCR-grade water and 2.5 µl of template. Cycling conditions were as follows: 94°C for 2 min, 35 amplification cycles (94°C for 30 s, 53°C for 45 s, 72°C for 1:30 min) and final extension at 72°C for 7 min. Considering that plant roots are colonised by multiple fungal species, PCR fragments were ligated into the pCR^®^4-TOPO^®^ vector and cloned into One Shot^®^ TOP10 competent *Escherichia coli* cells, following the manufacturer’s protocol for Kanamycin selection with modifications. Eight to 12 colonies per sample were directly amplified using ITS1F/ITS4 primers into 15 µl of PCR mix consisting of: 7.5 µl of JumpStart Taq ReadyMix, 0.075 µl of each primer (100 µM) and 7.35 µl of PCR-grade water. Cycling conditions were as follows: 94°C for 7 min, 25 amplification cycles (94°C for 30 s, 53°C for 45 s, 72°C for 1 min with the addition of 5 s per cycle during the extension phase), and final extension at 72°C for 7 min. Prior to sequencing, PCR products were purified using a 40% v/v dilution in PCR-grade water of a Exonuclease I : FastAP solution (ratio 1:2) (Thermo Fisher Scientific, Waltham, Massachusetts, United States). Purified PCR products were prepared for sequencing with BigDye v. 3.1 (Applied Biosystems,  Waltham, Massachusetts, United States) and primers ITS1F and ITS4.

Soil fungal communities

DNA was extracted in duplicate from 250 mg of soil (fresh weight) for each plot, using the DNeasy PowerSoil Pro kit (Qiagen, Hilden, Germany) according to manufacturer's instructions. DNA extracts were quantified using Quantus^TM^ Fluorometer with QuantiFluor® Dye System for dsDNA (Promega, Madison, Wisconsin, United States) and purity-checked (260/280nm and 260/230nm ratios) via NanoDrop spectrophotometer (Thermo Scientific, Waltham, Massachusetts, United States). DNA extracts were sent to Macrogen (Macrogen Genome Center, Seoul, Republic of Korea) for downstream processing and bioinformatic analysis. PCR amplification of the ITS2 region was performed using the primers ITS1F/ITS86F (White et al., 1990; Turenne et al., 1999), and amplicon libraries were sequenced on an Illumina MiSeq (Illumina, [San Diego, California, United States](https://www.google.com/search?sxsrf=ALiCzsbgjpTrJLUnGWAnyLJMkA4WYbB1pQ:1666106380138&q=San+Diego&stick=H4sIAAAAAAAAAOPgE-LSz9U3MDIvMUxPUeIAsc0Ny4q0tLKTrfTzi9IT8zKrEksy8_NQOFYZqYkphaWJRSWpRcWLWDmDE_MUXDJT0_N3sDLuYmfiYAAAOlk1YlgAAAA&sa=X&ved=2ahUKEwiBu8j3ier6AhWER8AKHZ0jB2wQmxMoAXoECHoQAw)). Reads pre-processing and clustering were carried out by Macrogen. Representative sequences from each OTU were assigned to a Species Hypothesis (SH) in house, using the SH matching v2.0.0 tool on PlutoF.

**Methods S4**. **Lichen photobiont analysis**

The lichen photobiont DNA was extracted using Chelex-100 resin (BioRad) following Ferencová et al. (2017). We analysed three specimens for each of the four lichen species found to be consistently present at each subplot. The ITS rDNA region was amplified with 7.5 μl of Green Taq Mastermix (Promega, Madison, WI), 0.7 μl of each primer at 10 μM concentration, and 1 μl of DNA (Pino-Bodas & Stenroos 2021). To determine to which *Asterochloris* lineage the photobionts sequenced belonged, the new sequences were included in the ITS rDNA alignment of Pino-Bodas & Stenroos (2021), with additional sequences from Kosecka et al. (2021), including the new lineages found by the authors. The alignment was made in MAFFT using default parameters (Katoh et al. 2019). The phylogeny was estimated with Maximum likelihood, implemented in RAxML 7.0.3 (Stamatakis et al. 2005) with GTRGAMMA model and 500 pseudoreplicates of rapid bootstrap. *Vulcanochloris canariensis* was used as outgroup. Because the taxonomy of the photobiont is still poorly understood (Muggia et al. 2018), most recent studies have used photobiont haplotypes to study the diversity and to infer the mycobiont-photobiont interaction (Pérez-Ortega et al. 2012; Cao et al. 2015; Garrido-Benavent et al. 2020, 2022). The number of photobiont haplotypes and genetic diversity in each plot was analysed with DnaSP (Librado & Rozas 2009). Haplotype networks under statistical parsimony were constructed in PopART 1.7 (Leigh & Bryant 2015) selecting the TCS method (Clement et al. 2000). Gaps were considered as missing data. Analysis of molecular variance (AMOVA) was conducted to determine the contribution of treatment in the genetic differentiation of photobionts, using ARLEQUIN V3.5 (Excoffier & Lischer 2010).
